# NAD⁺ modulation by nicotinamide riboside enhances CD8⁺ T-cell metabolic fitness and effector function during *Trypanosoma cruzi* infection and improves parasite control

**DOI:** 10.64898/2026.09.29.755262

**Authors:** Florencia Hellriegel, Camila Fontanari, Ruth E. Baigorri, Martín G. Theumer, Camila M. S. Giménez, Melisa Herrera, Luciana Mezzano, Claudia C. Motrán, Fabio M. Cerbán, Eva V. Acosta Rodriguez, Luciana Berod, Cinthia C. Stempin

## Abstract

Chagas disease, caused by the parasite *Trypanosoma cruzi*, remains a major health concern, with effective immune control relying heavily on CD8⁺ T cells. However, the metabolic factors that regulate CD8⁺ T cell function during infection are not well understood. Acute *T. cruzi* infection induced metabolic remodeling in parasite-specific and effector CD8⁺ T cells, characterized by increased reliance on glycolysis, altered mitochondrial homeostasis, oxidative stress, and increased CD38 expression. Nicotinamide riboside (NR), an NAD⁺ precursor, improved mitochondrial function, reduced mitochondrial reactive oxygen species (mROS) and CD38 expression, decreased apoptosis, and enhanced CD8⁺ T cell effector responses *in vitro* and in infected mice. *In vivo*, NR treatment increased intracellular NAD⁺ levels in CD8^+^ T cells and improved parasite control during acute infection, reducing parasitemia and skeletal muscle parasite burden. Furthermore, transient NR supplementation was associated with persistent metabolic and functional changes in memory CD8⁺ T cells during chronic infection. These findings identify NAD⁺ metabolism as a modulator of CD8⁺ T cell metabolic fitness during *T. cruzi* infection and support its further investigation as a host-directed strategy to enhance parasite control.

**Importance:** CD8⁺ T cells are essential for controlling *Trypanosoma cruzi*, the intracellular parasite that causes Chagas disease, yet the contribution of cellular metabolism to their protective function remains poorly understood. Using a murine model of *T. cruzi* infection, this study shows that metabolic and mitochondrial alterations accompany the CD8⁺ T cell response and that nicotinamide riboside supplementation enhances CD8⁺ T cell metabolic fitness and function while increasing intracellular NAD^+^ levels. These changes were associated with improved parasite control. Together, our findings highlight the metabolic state of CD8⁺ T cells as an important component of the immune response to *T. cruzi* and broaden our understanding of how host immunometabolism relates to the control of intracellular infection.

## Introduction

Chagas disease caused by the protozoan parasite *Trypanosoma cruzi* is endemic in Latin America although its prevalence continues to rise globally as a consequence of human migration (1). Disease progression, from an initial asymptomatic phase to severe cardiac and digestive pathologies, is shaped by parasite heterogeneity and by variability in the host immune response. CD8^+^ T cell mediated immunity is essential for parasite control throughout the various stages of infection (2, 3). CD8^+^ T cell deficient mice are extremely susceptible to infection (4, 5), and strategies that improve parasite-specific CD8^+^ T cell responses result in increased host protection (3, 6, 7). These findings, together with evidence that persistent parasitism correlates with disease severity in both human patients and experimental models (8–10), have fueled efforts to elucidate the factors governing effective CD8⁺ T cell-mediated immunity against *T. cruzi*. Accordingly, a deeper understanding of the mechanisms regulating CD8⁺ T cell activation, expansion, and effector function are critical to define their role in infection control and chronic disease progression.

In this context, cellular metabolism has emerged as a key determinant of T cell fate and function (11). Beyond energy production, metabolic programs actively regulate T cell activation, differentiation, and long-term function, shaping immune responses during infection and cancer (12). Although aerobic glycolysis supports the biosynthetic demands of rapid expanding effector CD8^+^ cells (13, 14), mitochondrial metabolism is also critical for sustaining T cell function (15–18).

A key metabolite that supports mitochondrial function is nicotinamide adenine dinucleotide (NAD^+^). NAD^+^ is a central cofactor that plays a pivotal role in cellular redox balance and energy metabolism including oxidative phosphorylation. Moreover, it serves as a substrate for several enzymes, such as sirtuin deacetylases (SIRTs), which regulate mitochondrial biogenesis and function in part through the peroxisome proliferator-activated receptor-γ coactivator 1α (PGC1α) (19–21). NAD^+^ metabolism has been shown to critically regulate mitochondrial fitness and T cell function, thereby shaping immune responses during infection, cancer, and aging (22–24). Thus, alterations in NAD^+^ homeostasis have been linked to impaired T cell responses (22, 25), while restoration of NAD^+^ levels through NAD-boosting strategies has been shown to limit CD8⁺ T cell exhaustion and promote effector differentiation in the tumor context (26, 27).

Intracellular NAD⁺ homeostasis depends on the balance between synthesis, degradation, and recycling, with the latter being the predominant pathway through vitamin B3 precursors such as nicotinamide, nicotinic acid, and nicotinamide riboside (NR) (28). NAD⁺ catabolism is primarily mediated by CD38, a transmembrane glycoprotein with NADase activity (29, 30). Inhibition of CD38 increases NAD⁺ levels (31), whereas supplementation with NR has been shown to increase NAD⁺ levels (32).

CD38 is expressed in activated T cells (33), and increased CD38 expression has been associated with metabolic and mitochondrial dysfunction, T cell exhaustion, and impaired effector function during infection and cancer (21, 22).

Here, we performed a comprehensive metabolic characterization of parasite-specific and effector CD8⁺ T cells in a murine model of *T. cruzi* infection and investigated whether increasing NAD⁺ availability with NR could influence CD8⁺ T cell metabolic fitness and function. We found that acute infection induces metabolic remodeling accompanied by increased CD38 expression and mitochondrial stress in parasite-specific and effector CD8⁺ T cells. NR supplementation increased intracellular NAD⁺ levels, improved mitochondrial fitness, enhanced CD8⁺ T cell effector responses, and was associated with improved parasite control.

## Results

### *T. cruzi* infection induces glycolytic reprogramming in parasite-specific CD8^+^ T cells

Sex-dependent differences in susceptibility and immune responses to *T. cruzi* infection have been reported, with male mice generally exhibiting greater susceptibility to acute infection (34, 35). Therefore, we initially characterized the metabolic remodeling of CD8⁺ T cells during *T. cruzi* infection in male mice. To this end, mice were infected intraperitoneally with 5,000 trypomastigotes (Tulahuen strain) and analyzed at different time points post-infection (0, 9, 20, and 78 dpi) (Fig. 1A). As expected, parasitemia peaked at 20 dpi and was reduced during the chronic phase of infection (Fig. 1B). Consistent with previous findings (36), the frequency of splenic TSKB20 tetramer-positive CD8⁺ T cells (hereafter referred to as parasite-specific CD8⁺ T cells, Tc-CD8⁺) increased during acute infection, reaching a peak at 20 dpi before undergoing contraction (Fig. 1C). A similar pattern was observed when Tc-CD8⁺ T cells were quantified as absolute numbers (Fig. S1A–C).

**Figure 1.**
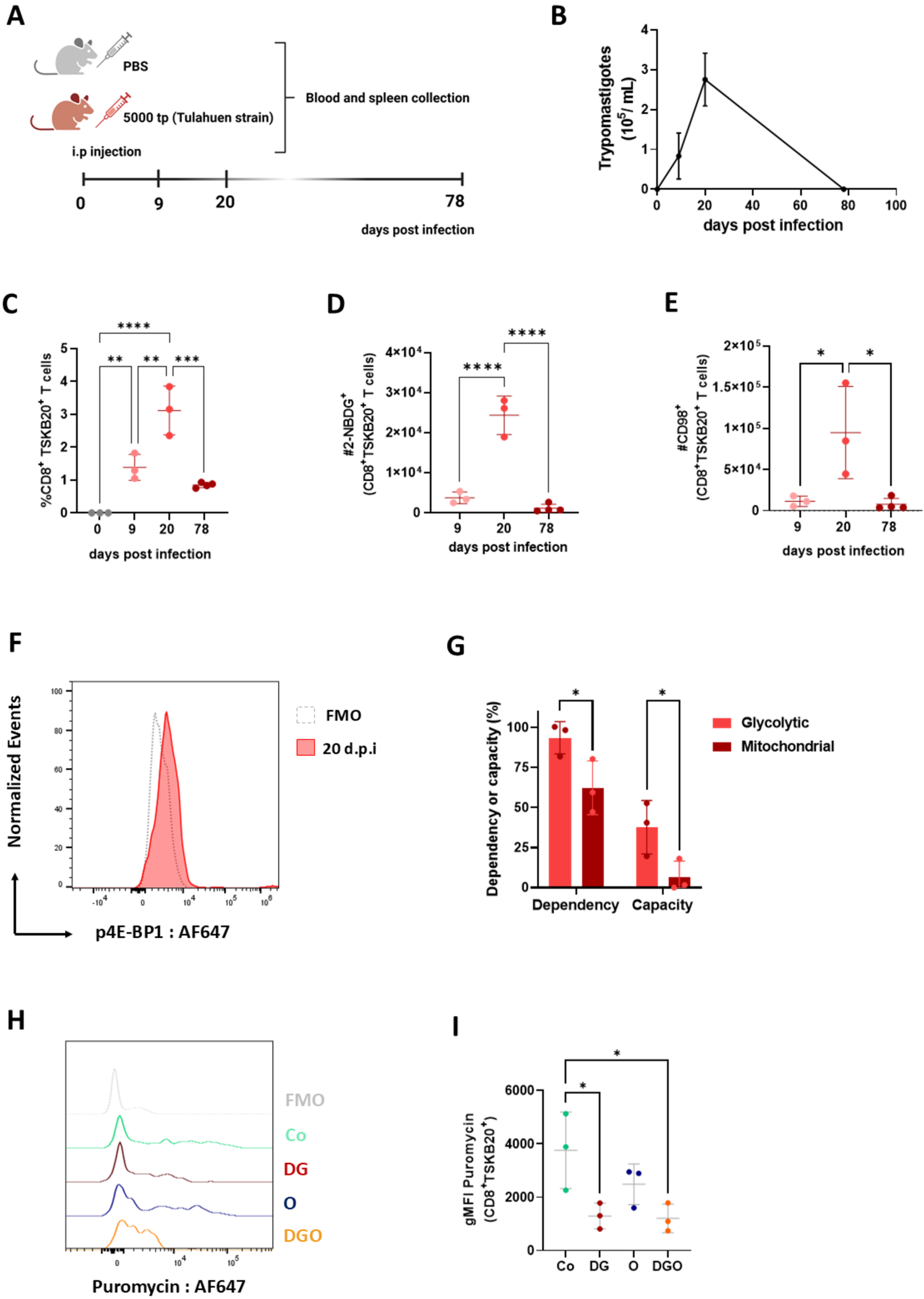
*T. cruzi* infection induces glycolytic reprogramming in parasite-specific CD8⁺ T cells. **(A)** Schematic representation of the experimental design. Male C57BL/6 mice were infected with 5,000 bloodstream trypomastigotes of the Tulahuen strain. Blood and spleen samples were collected at 9, 20 (acute phase, AP), and 78 days post-infection (dpi). **(B)** Blood parasitemia, expressed as the number of parasites per mL of blood, was determined by Neubauer chamber counting. **(C)** Relative frequency of CD8⁺ TSKB20^+^ T cells during the course of infection. **(D)** Glucose uptake was assessed using the fluorescent glucose analog 2-NBDG. Absolute numbers of CD8⁺ TSKB20^+^ 2-NBDG⁺ T cells are shown. **(E)** CD98 expression in CD8⁺ TSKB20^+^ T cells was evaluated by flow cytometry. Absolute numbers are shown. **(F)** Levels of Phosphorylated 4E-BP1 (p-4E-BP1) in parasite-specific CD8⁺ T cells (TSKB20^+^) were analyzed by flow cytometry. A representative histogram and the corresponding fluorescence minus one (FMO) control are shown. **(G)** SCENITH analysis was performed on splenocytes isolated from AP mice and stimulated with *T. cruzi* lysate for 18 h. Bar graphs show glycolytic dependence, mitochondrial metabolic dependence, glycolytic capacity, and mitochondrial metabolic capacity of TSKB20^+^ CD8⁺ T cells. **(H)** Representative histograms of puromycin incorporation under each SCENITH condition: control (Co), 2-deoxy-D-glucose (DG), oligomycin (O), and a combination of both (DGO). **(I)** Mean fluorescence intensity (MFI) of puromycin under each experimental condition. Data were obtained from 9 dpi (*n* = 3), 20 dpi (*n* = 3), and 78 dpi (*n* = 4) mice per group and are presented as mean ± SD. Data in (C), (D), (E), and (I) were analyzed using one-way ANOVA followed by Tukey’s multiple-comparisons test. Data in (G) were analyzed using two-way ANOVA followed by Šídák’s multiple-comparisons test. ns, not significant; \**p* < 0.05, \*\**p* < 0.01, \*\*\**p* < 0.001, and \*\*\*\**p* < 0.0001.

To characterize the metabolic status of Tc-CD8⁺ T cells, we first evaluated glucose uptake and amino acid transport by measuring 2-NBDG uptake and CD98 expression, respectively. The absolute numbers of 2-NBDG⁺ and CD98⁺ Tc-CD8⁺ T cells increased during the acute phase of infection (20 dpi) (Fig. 1D-E). CD98 expression was also elevated, as shown by representative histograms and increased mean fluorescence intensity (MFI) (Fig. S1D-E). Given that increased glucose uptake and amino acid availability are consistent with activation of the mTOR pathway, we next assessed mTOR activation at 20 dpi by measuring phosphorylation of 4E-BP1, a downstream target of mTORC1. Tc-CD8⁺ T cells showed detectable p-4E-BP1 levels above the fluorescence minus one (FMO) control, consistent with activation of the mTOR pathway during the acute phase of infection (Fig. 1F).

We next evaluated the metabolic profile of Tc-CD8⁺ T cells using SCENITH (<u>S</u>ingle <u>C</u>ell <u>EN</u>ergetIc metabolism by profiling <u>T</u>ranslation in<u>H</u>ibition) (37). This approach estimates metabolic dependencies and capacities at the single-cell level by measuring puromycin incorporation as a readout of protein synthesis following inhibition of glucose metabolism with 2-deoxyglucose (2-DG) and mitochondrial ATP production with oligomycin (O). Splenocytes from infected mice were collected at 20 dpi, stimulated with *T. cruzi* lysate, and subjected to SCENITH analysis. Tc-CD8⁺ T cells exhibited high glycolytic dependency together with reduced mitochondrial metabolic capacity (Fig. 1G-I). These findings indicate that parasite-specific CD8⁺ T cells preferentially rely on glycolysis and have reduced mitochondrial metabolic capacity during acute infection.

### Mitochondrial stress is increased in CD38^hi^ parasite-specific CD8⁺ T cells during acute *T. cruzi* infection

To further characterize the mitochondrial phenotype of Tc-CD8⁺ T cells during acute infection, we evaluated multiple parameters of mitochondrial function by flow cytometry. Mitochondrial ROS (mROS) was assessed using MitoSOX, mitochondrial mass (MM) was measured with MitoSpy™ Green (MT Green; MTg), and mitochondrial membrane potential (MMP) was evaluated using MitoStatus Red (MTst Red; MTst). Analysis of Tc-CD8⁺ T cells revealed an increased frequency of mROS⁺ cells together with elevated mROS levels at 9 and 20 dpi, peaking during the acute phase (Fig. 2A–B). In addition, Tc-CD8⁺ T cells exhibited increased MM and reduced MMP during the acute phase (Fig. 2C-D), with an accumulation of MTst^−^MTg^+^ cells, consistent with mitochondrial depolarization and increased mitochondrial stress (Fig. 2E-F). These alterations were attenuated by 78 dpi.

**Figure 2.**
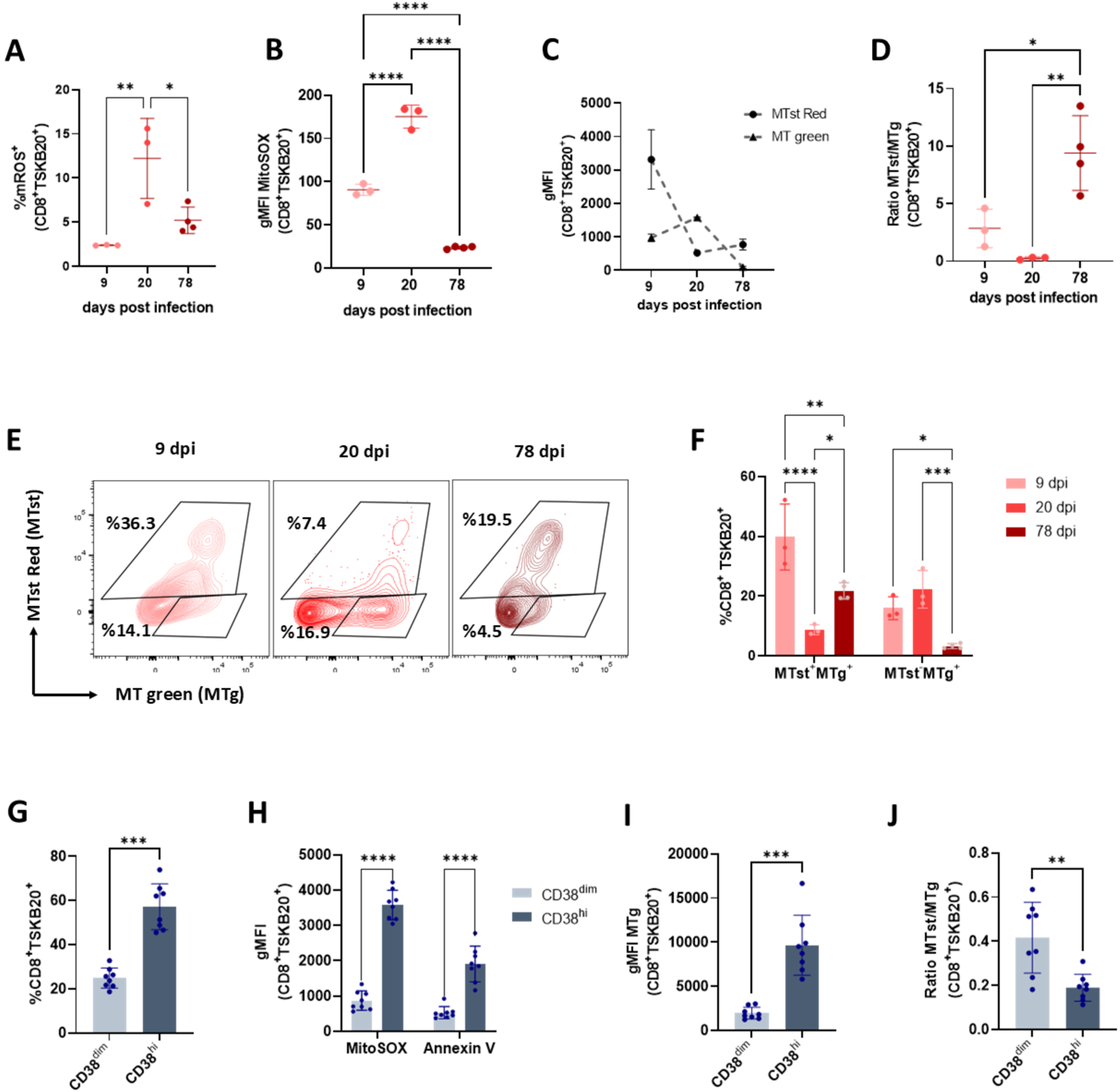
Mitochondrial stress is increased in CD38^hi^ parasite-specific CD8⁺ T cells during acute *T. cruzi* infection. **(A-B)** *Ex vivo* mitochondrial reactive oxygen species (mROS) levels were analyzed in TSKB20^+^ CD8⁺ T cells from infected male mice at 9, 20, and 78 days post-infection (dpi) using MitoSOX. **(A)** Frequency of mROS^+^ cells. **(B)** Mean fluorescence intensity (MFI) of MitoSOX. **(C-F)** Mitochondrial parameters were analyzed in TSKB20^+^ CD8⁺ T cells at 9, 20, and 78 dpi by flow cytometry using MitoSpy™ Green (MTg), which labels mitochondria independently of mitochondrial membrane potential, together with the membrane potential-dependent probe MitoStatus Red (MTst). These probes were used to identify cells with preserved mitochondrial membrane potential (MTst⁺MTg⁺) or depolarized mitochondria (MTst⁻MTg^+^) **(C)** MFI of MTg and MTst at the indicated time points of infection. **(D)** MTg/MTst ratio. **(E)** Representative contour plots showing TSKB20^+^ CD8⁺ T cells with preserved (MTst⁺MTg⁺) or depolarized (MTst⁻MTg^+^) mitochondria at each time point. **(F)** Percentage of MTst⁺MTg⁺ and MTst⁻MTg^+^ TSKB20^+^ CD8⁺ T cells. **(G–J)** CD38 expression was analyzed in TSKB20^+^ CD8⁺ T cells at 20 dpi. **(G)** Frequencies of CD38^dim^ and CD38^hi^ subsets within TSKB20⁺ CD8⁺ T cells. **(H)** MFI of MitoSOX and Annexin V in CD38^dim^ and CD38^hi^ populations. **(I)** MFI of MitoSpy™ Green (MTg) and **(J)** MTg/MTst ratio in CD38^dim^ and CD38^hi^ populations. Data for panels (A–F) were obtained from mice at 9 dpi (*n* = 3), 20 dpi (*n* = 3), and 78 dpi (*n* = 4). Data for panels (G–J) were obtained from mice at 20 dpi (*n* = 8). Data are presented as mean ± SD. Data in (A), (B), and (D) were analyzed using one-way ANOVA followed by Tukey’s multiple-comparisons test. Data in (F) and (H) were analyzed using two-way ANOVA followed by Šídák’s multiple-comparisons test. Data in (G) and (I-J) were analyzed using a paired two-tailed Student’s t-test. \**p* < 0.05, \*\**p* < 0.01, \*\*\**p* < 0.001, and \*\*\*\**p* < 0.0001.

Given that mitochondrial alterations were most pronounced during the acute phase, we next focused our analysis on Tc-CD8⁺ T cells at 20 dpi. Considering the role of CD38 in NAD⁺ metabolism, we next evaluated CD38 expression in Tc-CD8⁺ T cells and stratified them into CD38^hi^ and CD38^dim^ subsets for subsequent analyses. During the acute phase of infection, the frequency of CD38^hi^ Tc-CD8⁺ T cells was higher than that of the CD38^dim^ subset (Fig. 2G, Fig. S1F). Notably, CD38^hi^ Tc-CD8⁺ T cells displayed higher mROS levels, increased Annexin V staining, elevated mitochondrial mass, and reduced mitochondrial membrane potential compared with the CD38^dim^ subset (Fig. 2H-J).

Together, these results indicate that CD38^hi^ parasite-specific CD8 T cells exhibit increased oxidative stress, apoptosis, and mitochondrial alterations during acute *T. cruzi* infection.

### High CD38 expression characterizes stressed effector CD8⁺ T cells during acute *T. cruzi* infection

Because the low frequency of Tc-CD8⁺ T cells limited more detailed phenotypic analyses, we extended our study to the total effector CD8⁺ T cell compartment. As expected, acute infection induced a marked expansion of effector CD8⁺ T cells (CD44⁺CD62L^low^) compared with non-infected animals (Fig. S2A–B). Within this population, t-SNE analysis identified clusters of CD44⁺CD62L^low^ CD8⁺ T cells characterized by increased PD-1 and CD38 expression, reduced MMP, and elevated mROS levels, consistent with the phenotype observed in Tc-CD8⁺ T cells (Fig. 3A). Moreover, SCENITH analysis showed that effector CD8⁺ T cells exhibited a metabolic profile characterized by high glycolytic dependency and reduced mitochondrial capacity (Fig. S2C-F).

**Figure 3.**
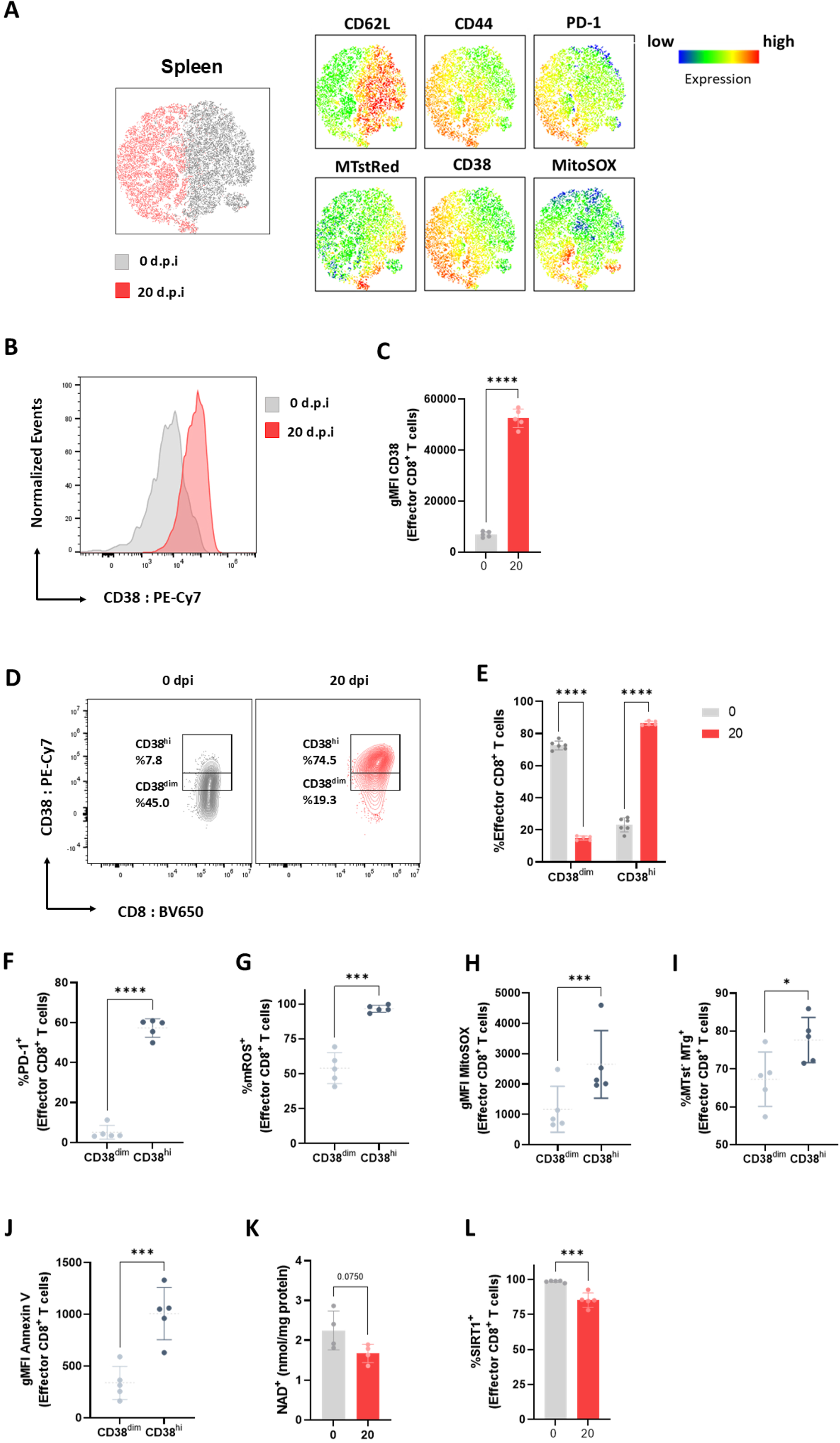
High CD38 expression characterizes stressed effector CD8⁺ T cells during acute *T. cruzi* infection. Splenic CD8⁺ T cells from non-infected (0 dpi) and acutely infected (20 dpi) male mice were analyzed by flow cytometry. **(A)** Representative t-distributed stochastic neighbor embedding (t-SNE) analysis of splenic CD8⁺ T cells. Expression of CD62L, CD44, PD-1, MitoStatus Red (MTst), CD38, and MitoSOX is displayed as heat maps on the t-SNE projection. **(B-J)** Analyses were performed on effector CD8⁺ T cells (CD44⁺CD62L^low^). **(B)** Representative histograms of CD38 expression. **(C)** Mean fluorescence intensity (MFI) of CD38. **(D)** Representative contour plots showing CD38^dim^ and CD38^hi^ effector CD8⁺ T cell populations. **(E-J)** Effector CD8⁺ T cells were stratified into CD38^dim^ and CD38^hi^ subsets for subsequent analyses. **(E)** Frequencies of CD38^dim^ and CD38^hi^ effector CD8⁺ T cells. **(F)** Frequency of PD-1⁺ cells. **(G)** Frequency of mROS⁺ cells. **(H)** Mean fluorescence intensity (MFI) of MitoSOX. **(I)** Frequency of cells with depolarized mitochondria (MTst⁻MTg⁺). **(J)** Mean fluorescence intensity (MFI) of Annexin V. **(K)** Intracellular NAD⁺ levels in purified bulk CD8⁺ T cells. (**L**) Frequency of SIRT1⁺ effector CD8⁺ T cells. Data were obtained from 0 dpi (*n* = 4) and 20 dpi (*n* = 5) mice per group. For panel (L) data were obtained from 0 dpi (*n* = 4) and 20 dpi (*n* = 4) mice per group. Data in (C), (K), and (L) were analyzed using an unpaired two-tailed Student’s t-test. Data in (F–J) were analyzed using a paired two-tailed Student’s t-test. Data in (E) were analyzed using two-way ANOVA followed by Šídák’s multiple-comparisons test. \**p* < 0.05, \*\*\**p* < 0.001, and \*\*\*\**p* < 0.0001.

We next evaluated CD38 expression within the effector CD8⁺ T cell compartment. Effector CD8⁺ T cells from infected mice exhibited increased CD38 expression compared with non-infected controls (Fig. 3B-C). Given the marked increase in CD38 expression, effector CD8⁺ T cells were further stratified according to CD38 expression levels into CD38^hi^ and CD38^dim^ subsets. Infection led to a shift in the distribution of CD38 expression, with an increased frequency of CD38^hi^ cells and a concomitant reduction in CD38^dim^ cells (Fig. 3D-E). We then directly compared CD38^hi^ and CD38^dim^ effector CD8⁺ T cells during infection. CD38^hi^ cells displayed increased frequencies of PD-1^+^ and mROS^+^ cells, higher mROS levels, loss of MMP and increased Annexin V staining (Fig. 3F-J), consistent with the phenotype observed in Tc-CD8⁺ T cells.

Given the NAD⁺-consuming activity of CD38, we next quantified intracellular NAD⁺ levels in purified CD8⁺ T cells from infected and non-infected mice. CD8⁺ T cells from infected mice showed a trend toward lower intracellular NAD⁺ levels compared with cells from non-infected controls (Fig. 3K). We also evaluated SIRT1, an NAD⁺-dependent deacetylase, and found that the frequency of SIRT1⁺ effector CD8⁺ T cells was significantly reduced during infection (Fig. 3L, Fig. S2G).

Together, these findings support an association between increased CD38 expression and metabolic and mitochondrial alterations observed in effector CD8⁺ T cells during acute infection.

### Nicotinamide riboside modulates proliferation and CD38 expression during effector CD8⁺ T cell differentiation *in vitro*

To determine whether modulation of NAD metabolism could directly influence CD8⁺ T cell function, we established an *in vitro* differentiation system using naïve CD8⁺ T cells. Cells were activated and differentiated into effector CD8⁺ T cells in the presence or absence of nicotinamide riboside (NR), a precursor of NAD⁺, thereby allowing us to assess the direct effects of NR on CD8^+^ T cells independently of the infection context (Fig. 4A). We first assessed whether NR affected cell survival. No significant differences in cell viability were observed between NR-treated and control conditions (Fig. S3A-B), indicating that NR does not impair CD8⁺ T cell survival during differentiation.

**Figure 4.**
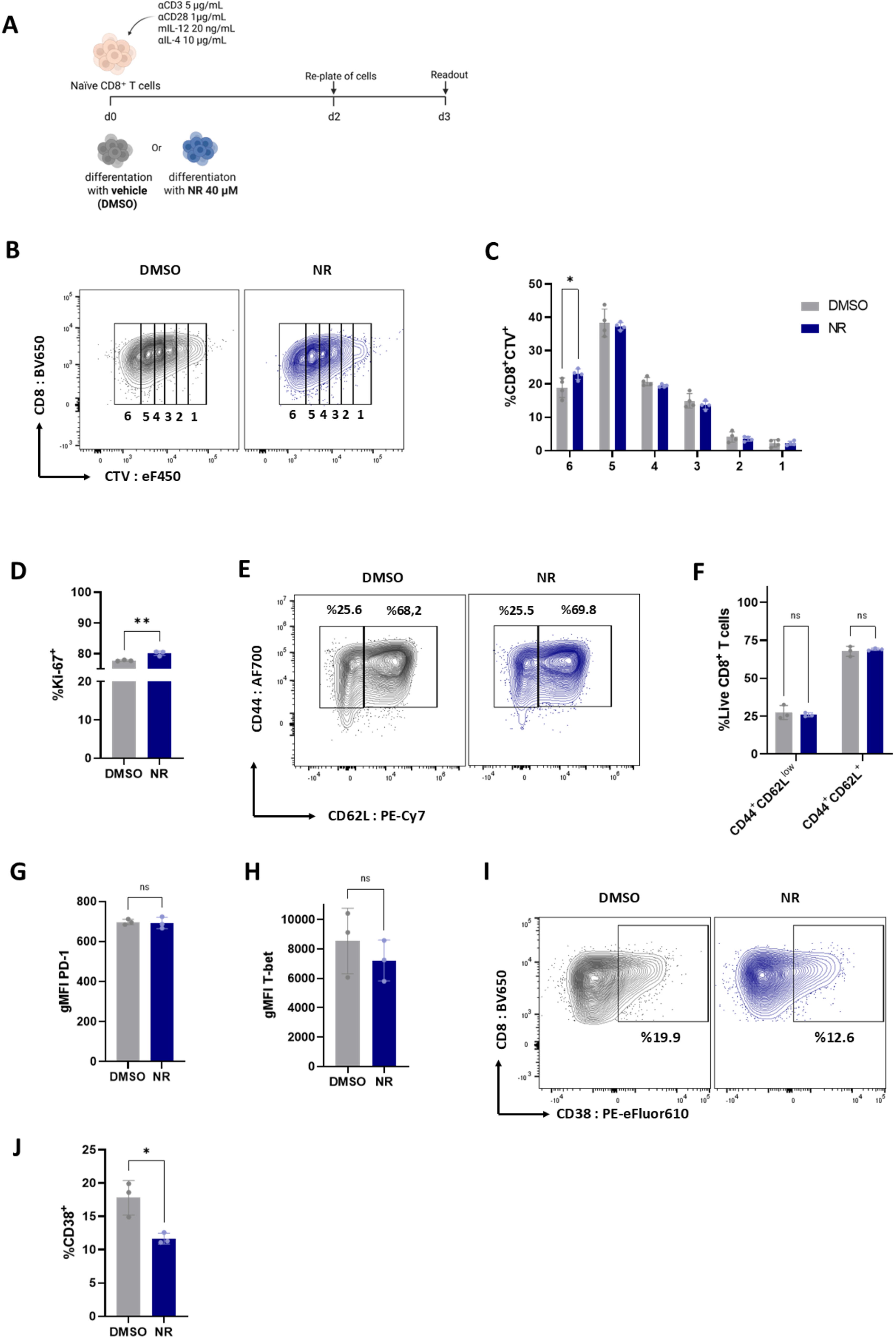
Nicotinamide riboside modulates proliferation and CD38 expression during effector CD8⁺ T cell differentiation *in vitro*. For each independent experiment, naïve CD8⁺ T cells isolated from C57BL/6 mice (*n =*4) were pooled and cultured under effector-polarizing conditions in the presence of DMSO (vehicle) or 40 µM nicotinamide riboside (NR). **(A)** Schematic representation of the experimental design. **(B-C)** Cell proliferation was assessed by CellTrace™ Violet (CTV) dilution. **(B)** Representative contour plots showing CTV-labeled live CD8⁺ T cells. **(C)** Frequency of CTV⁺ CD8⁺ T cells at the indicated division rounds. **(D-H)** Differentiation and activation status were analyzed by flow cytometry. **(D)** Frequency of Ki-67⁺ CD8⁺ T cells. **(E)** Representative contour plots showing CD44⁺CD62L⁺ and CD44⁺CD62L^low^ CD8⁺ T cell populations. **(F)** Frequencies of CD44⁺CD62L⁺ and CD44⁺CD62L^low^ CD8⁺ T cells. **(G)** Mean fluorescence intensity (MFI) of PD-1. **(H)** MFI of T-bet. **(I, J)** CD38 expression was analyzed by flow cytometry. **(I)** Representative contour plots of CD38 expression. **(J)** Frequency of CD38⁺ CD8⁺ T cells. Data are presented as mean ± SD of three to four technical replicates from one experiment representative of three independent experiments. Data in (C) and (F) were analyzed using two-way ANOVA followed by Šídák’s multiple-comparisons test. Data in (D), (G), (H), and (J) were analyzed using an unpaired two-tailed Student’s t-test. ns, not significant; \**p* < 0.05, \*\**p* < 0.01.

We next evaluated cell proliferation using CellTrace dilution and found an increase in the proportion of NR-treated cells reaching the sixth division peak compared with control cells (Fig. 4B-C). Consistently, although the majority of cells were Ki-67⁺ under both conditions, NR-treated CD8⁺ T cells exhibited a significant increase in the frequency of Ki-67⁺ cells (Fig. 4D), further supporting an effect of NR on CD8⁺ T cell proliferation. We next analyzed key markers of effector differentiation. The frequencies of CD44⁺CD62L^low^ and CD44⁺CD62L⁺ CD8⁺ T cells were comparable between NR-treated and control cultures (Fig. 4E-F). Likewise, no significant differences were observed in the expression levels of PD-1⁺ or T-bet⁺ cells between groups (Fig. 4G-H), indicating that NR does not alter the acquisition of the effector phenotype. Despite the absence of changes in effector differentiation, NR treatment resulted in a reduced frequency of CD38⁺ cells compared with control conditions (Fig. 4I-J), suggesting that NR modulates CD38 expression during effector differentiation.

### Nicotinamide riboside enhances mitochondrial fitness and favors cytokine production by CD8⁺ T cells *in vitro*

To evaluate whether NR modulates cellular metabolism and mitochondrial function, we performed Seahorse extracellular flux analysis. NR-treated CD8⁺ T cells exhibited increased basal and maximal mitochondrial respiration, as well as increased ATP production, compared with control cells (Fig. 5A-B). In contrast, no significant differences were observed in extracellular acidification rate (ECAR) (Fig. S3C-D), indicating that NR enhanced mitochondrial oxidative metabolism without significantly affecting glycolytic activity under these experimental conditions. Consistent with these findings, NR-treated cells showed increased mRNA expression of *Sirt1* (Fig. 5C) and *Ppargc1a* (Fig. 5D), key regulators of mitochondrial function and biogenesis. In line with improved mitochondrial function, NR-treated CD8⁺ T cells displayed a reduced proportion of cells with depolarized mitochondria, indicating preservation of MMP (Fig. 5E-F).

**Figure 5.**
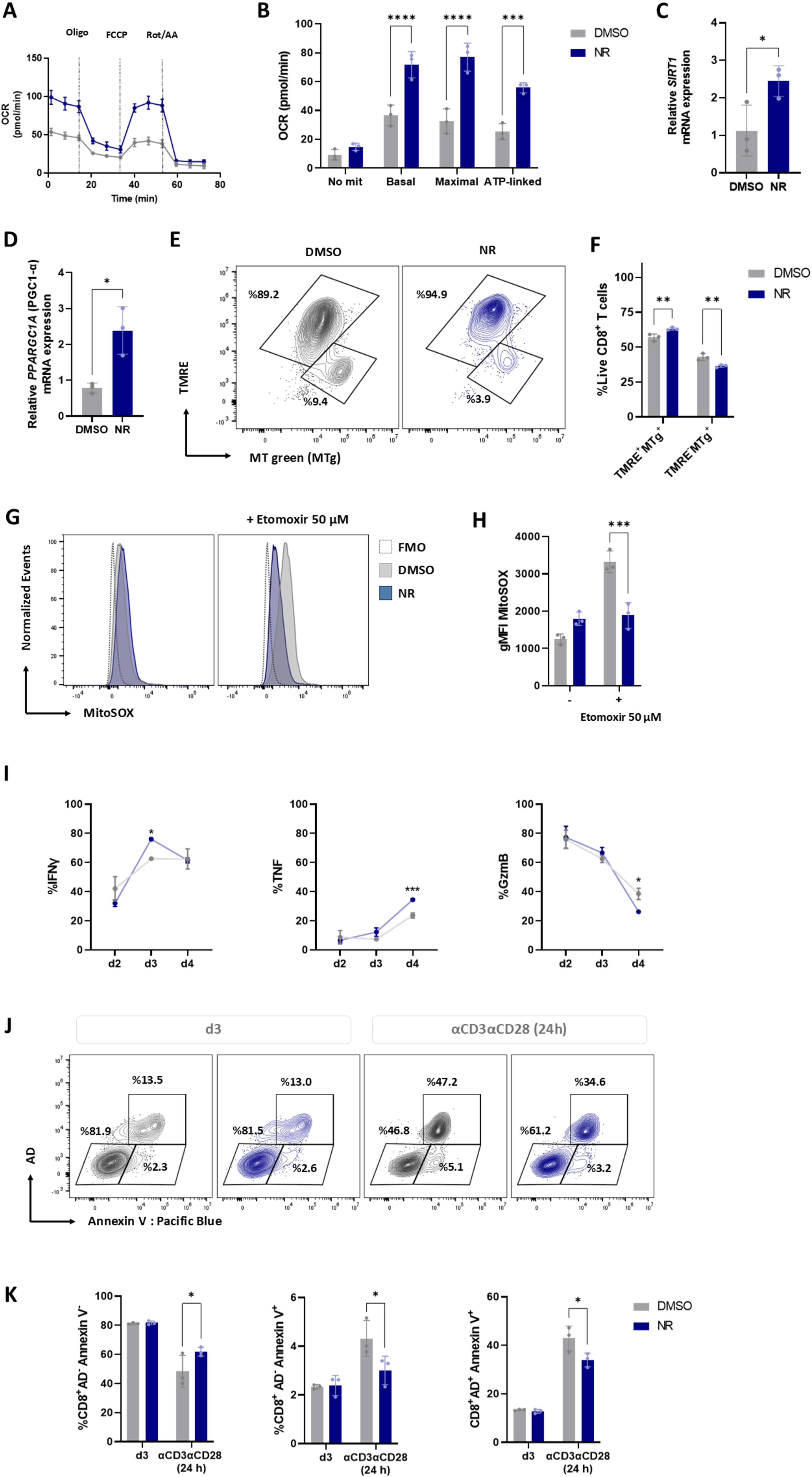
Nicotinamide riboside enhances mitochondrial fitness and favors cytokine production by CD8⁺ T cells *in vitro.* For each independent experiment, naïve CD8⁺ T cells were obtained from C57BL/6 mice (*n* = 4) and differentiated under effector-polarizing conditions in the presence of DMSO (vehicle) or 40 µM nicotinamide riboside (NR). **(A-B)** Mitochondrial respiration was evaluated using the Seahorse XF Cell Mito Stress Test. **(A)** Representative oxygen consumption rate (OCR) profile following sequential injection of oligomycin, FCCP, and rotenone/antimycin A. **(B)** Quantification of non-mitochondrial respiration, basal respiration, maximal respiration, and ATP-linked respiration. **(C)** Relative mRNA expression of *Sirt1* and **(D)** *Ppargc1a*. **(E-F)** MitoSpy™ Green (MTg), which labels mitochondria independently of mitochondrial membrane potential, together with tetramethylrhodamine ethyl ester (TMRE), a membrane potential-dependent probe, was used to identify cells with preserved mitochondrial membrane potential (TMRE⁺MTg⁺) or depolarized mitochondria (TMRE⁻MTg⁺). **(E)** Representative contour plots. **(F)** Frequencies of cells with preserved mitochondrial membrane potential (TMRE⁺MTg⁺) and cells with depolarized mitochondria (TMRE⁻MTg⁺). **(G-H)** Cells were treated with etomoxir (50 μM) for 15 min or left untreated, and mitochondrial reactive oxygen species (mROS) were assessed using MitoSOX. **(G)** Representative histograms of MitoSOX fluorescence. **(H)** Mean fluorescence intensity (MFI) of MitoSOX. **(I)** Intracellular production of IFN-γ, TNF, and granzyme B was evaluated by flow cytometry on days 2, 3, and 4 of culture. Frequencies of IFN-γ, TNF and granzyme B-producing CD8⁺ T cells are shown. **(J-K)** Apoptosis was assessed in CD8⁺ T cells on day 3 of culture and after 24 h of restimulation with plate-bound anti-CD3 (5 μg/mL) and soluble anti-CD28 (1 μg/mL). **(J)** Representative contour plots. **(K)** Frequencies of Aqua Dead dye (AD)⁺Annexin V⁺, AD⁺Annexin V⁻, and AD⁻Annexin V⁺ cells. Data are presented as mean ± SD. Data in (A) and (B) are representative of two independent experiments, whereas data in the remaining panels are representative of three independent experiments. Data are presented as mean ± SD. Data in (B), (F), (H), (I), and (K) were analyzed using two-way ANOVA followed by Šídák’s multiple-comparisons test. Data in (C) and (D) were analyzed using an unpaired two-tailed Student’s t-test. \**p* < 0.05, \*\**p* < 0.01, \*\*\**p* < 0.001, and \*\*\*\**p* < 0.0001.

To evaluate the ability of NR-treated CD8⁺ T cells to withstand mitochondrial stress, differentiated cells were exposed to a high concentration of etomoxir, a condition previously reported to induce mitochondrial dysfunction and oxidative stress beyond CPT1 inhibition (38). Following etomoxir treatment, NR-differentiated cells exhibited markedly reduced mROS levels (Fig. 5G-H), together with a lower frequency of mROS⁺ cells (Fig. S3E), while cell viability remained unchanged (Fig. S3F-G), indicating enhanced resistance to mitochondrial oxidative stress.

Given the impact of NR on mitochondrial fitness, we next assessed its effects on CD8⁺ T cell effector function. NR-differentiated CD8⁺ T cells exhibited a significant increase in the frequency of IFN-γ^+^ cells at day 3 and TNF^+^ cells at day 4 compared with DMSO-treated controls. In contrast, the frequency of granzyme B^+^ cells was reduced at day 4, indicating that NR differentially modulates CD8^+^ effector functions (Fig. 5I). To determine whether these effects were maintained following secondary stimulation, effector-differentiated CD8⁺ T cells were subjected to polyclonal restimulation (Fig. S4A). Upon restimulation, NR-treated cells displayed enhanced IFN-γ and TNF production (Fig. S4B-F). In addition, cells differentiated in the presence of NR exhibited an increased frequency of viable cells (AD⁻Annexin V⁻) and lower frequencies of early (AD⁻Annexin V⁺) and late apoptotic cells (AD⁺Annexin V⁺) compared with controls (Fig. 5J-K).

Together, these results demonstrate that NR enhances mitochondrial function, reduces oxidative stress and modulates CD8⁺ T cell effector responses, favoring IFN-γ and TNF production.

### NR improves parasite control and enhances CD8⁺ T cell responses during acute *T. cruzi* infection

Given the beneficial effects of NR on CD8⁺ T cell mitochondrial fitness and function *in vitro*, we next investigated whether NR supplementation could enhance CD8⁺ T cell responses and parasite control during *T. cruzi* infection *in vivo*. Considering the reported sexual dimorphism in susceptibility and immune responses to *T. cruzi* infection (34, 35), we first examined whether CD38 expression differed between male and female mice. Despite these known sex-related differences, effector CD8⁺ T cells from infected males and females displayed comparable levels of CD38 expression at 20 dpi (Fig. S5A). We then evaluated NR supplementation in both sexes, which were analyzed separately. To this end, infected mice were treated daily with nicotinamide riboside (NR, 500 mg/kg/day) or PBS from day 5 to day 20 dpi and analyzed at 21 dpi during the acute phase of infection (Fig. 6A). Age-matched non-infected mice received the corresponding treatment following the same schedule. We first examined the effects of NR supplementation in male mice. NR treatment improved control of infection, as evidenced by a reduction in parasitemia in treated mice compared with PBS controls (Fig. 6B). This effect was accompanied by a marked decrease in tissue parasite burden in skeletal muscle (SM), whereas no significant differences were observed in the heart or liver (Fig. 6C). Consistent with the lower SM parasite burden detected by qPCR during acute infection, histopathological analysis revealed fewer amastigote nests in NR-treated mice (Fig. 6D-E).

**Figure 6.**
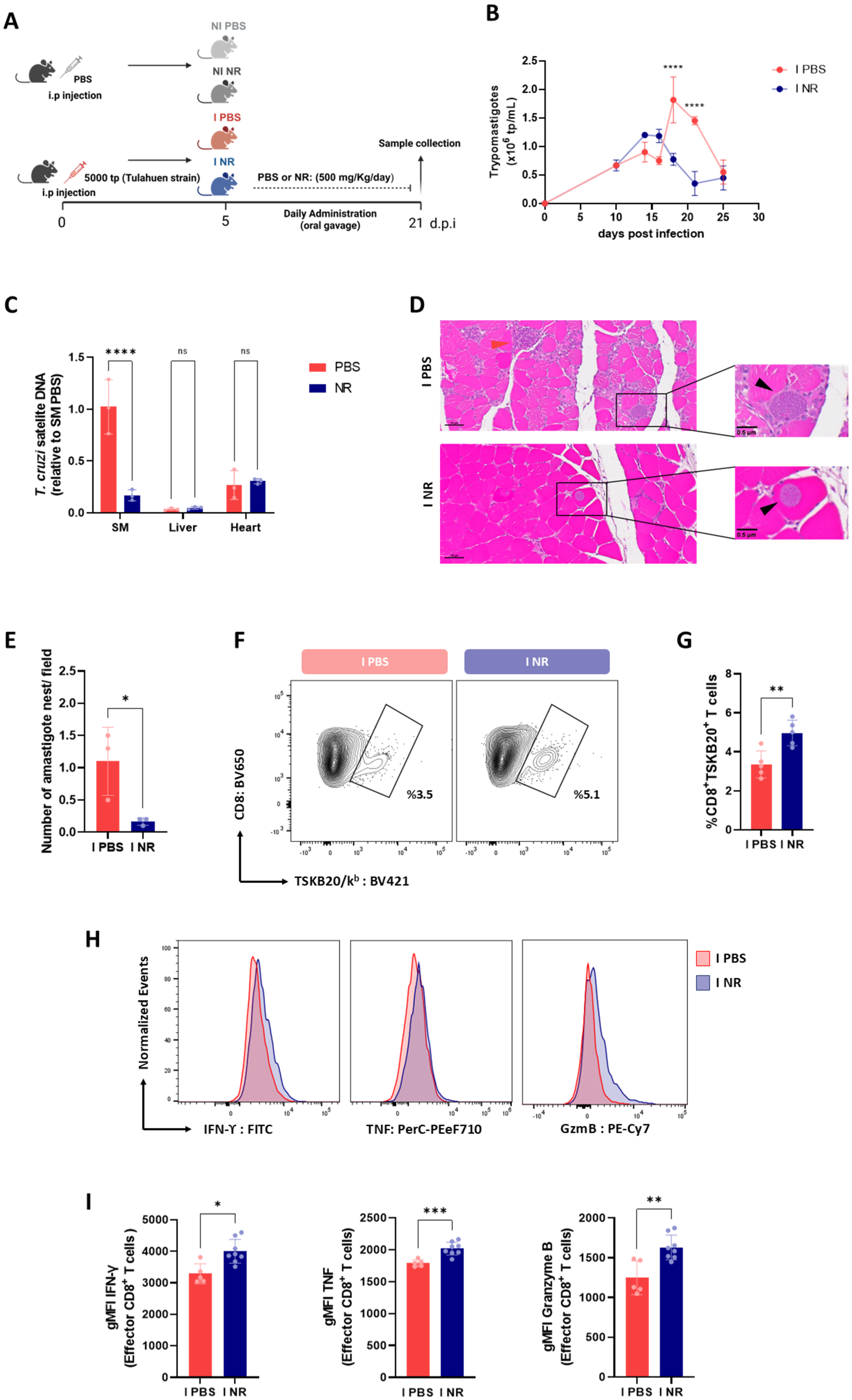
NR improves parasite control and enhances CD8⁺ T cell responses during acute *T. cruzi* infection. Male C57BL/6 mice were infected with 5,000 bloodstream trypomastigotes and treated daily with nicotinamide riboside (NR; 500 mg/kg) or PBS by oral gavage from 5 to 20 days post-infection (dpi). Age-matched non-infected mice receiving PBS or NR were included as controls (NI PBS and NI NR). **(A)** Schematic representation of the experimental design. **(B)** Blood parasitemia was monitored from 0 to 25 dpi and expressed as the number of parasites per mL of blood. **(C)** Tissue parasite burden was quantified by real-time PCR in pooled skeletal muscle (SM), liver, and heart from I PBS (*n* = 5) and I NR (*n* = 7) mice and expressed relative to SM of I PBS mice. **(D-E)** Histopathological analysis of amastigote nests in SM. **(D)** Representative images of SM sections from PBS- and NR-treated infected mice. Black arrows indicate amastigote nests, whereas red arrows identify nests surrounded by inflammatory infiltrates. Scale 50 µm. **(E)** Numbers of amastigote nests were quantified in ten fields per sample. **(F-G)** Parasite-specific CD8⁺ T cells were identified by flow cytometry using TSKB20^+^ tetramers. **(F)** Representative contour plots. **(G)** Frequency of TSKB20^+^ CD8⁺ T cells. **(H-I)** Effector response was evaluated in CD8⁺ T cells (CD44⁺CD62L^low^) by flow cytometry following *ex vivo* restimulation with PMA/ionomycin. **(H)** Representative histograms of IFN-γ, TNF, and granzyme B expression. **(I)** Mean fluorescence intensity (MFI) of IFN-γ, TNF, and granzyme B. Data were obtained from I PBS (*n* = 5) and I NR (*n* = 5) mice for panels B and G; from I PBS (*n* =3) and I NR (*n* =3) for panel E and from I PBS (*n* = 5) and I NR (*n* = 8) mice for panel I. For panel (C), each symbol represents one technical replicate. Data are presented as mean ± SD. Data in (B) and (C) were analyzed using two-way ANOVA followed by Šídák’s multiple-comparisons test. Data in (E), (G), and (I) were analyzed using an unpaired two-tailed Student’s t-test. ns, not significant; \**p* < 0.05, \*\**p* < 0.01, \*\*\**p* < 0.001, and \*\*\*\**p* < 0.0001.

We next investigated whether the improved control of infection observed in NR-treated mice was associated with changes in the CD8⁺ T cell response. While NR treatment did not significantly alter the overall distribution of CD8⁺ T cell subsets (Fig. S5B-C), it resulted in an increased frequency of Tc-CD8⁺ T cells (Fig. 6F-G).

Functionally, CD8⁺ T cells from NR-treated mice exhibited enhanced effector responses, including an increased frequency of IFN-γ⁺ cells (Fig. S5D-E) and higher IFN-γ, TNF, and granzyme B expression levels, compared with PBS-treated controls (Fig. 6H-I). Similarly, splenocytes from NR-treated mice produced higher levels of IFN-γ following *ex vivo* restimulation (Fig. S5F).

To determine whether the reduced parasite burden observed in NR-treated mice was accompanied by histopathological changes, SM and liver sections from NR-treated and PBS-treated infected mice, as well as non-infected controls, were examined. SM from infected mice exhibited inflammatory infiltrates, alterations in muscle architecture, and myofiber necrosis. Although NR-treated mice tended to exhibit milder histopathological involvement than PBS-treated mice, the severity of these alterations varied among individual animals (Fig. S6A–B). Liver sections from infected mice showed mild inflammatory infiltrates, steatosis, vascular congestion, and alterations in lobular architecture. These histopathological features were comparable between PBS- and NR-treated infected animals. However, fibrosis was detected only in PBS-treated animals and was absent in NR-treated animals (Fig. S6C-D).

We next assessed whether the improved parasite control observed in NR-treated mice was accompanied by changes in biochemical parameters indicative of disease severity. Infection induced a marked increase in serum LDH and CPK levels compared with non-infected animals (Fig. S6E), indicating tissue injury. In addition, infected mice exhibited reduced blood glucose levels relative to non-infected controls (Fig. S6F). However, NR treatment did not significantly change LDH, CPK, or blood glucose levels in infected mice compared with PBS-treated infected controls (Fig. S6E-F). Likewise, body weight remained comparable between infected groups (Fig. S6G). Importantly, NR administration to non-infected mice did not affect body weight or serum markers of tissue damage.

Together, these findings indicate that in male mice, NR supplementation enhances CD8⁺ T cell immunity and improves parasite control during acute *T. cruzi* infection without exacerbating systemic markers of tissue injury or disease severity.

### NR reduces CD38 expression and metabolic stress in CD8⁺ T cells during acute infection

Given the association between CD38 expression and mitochondrial dysfunction observed during infection, we next evaluated whether NR treatment modulated the metabolic profile of CD8⁺ T cells in male mice *in vivo*.

NR-treated mice exhibited reduced CD38 expression levels in effector CD8⁺ T cells compared with controls (Fig. 7A-B). Consistently, NR treatment decreased the frequency of CD38⁺ effector CD8⁺ T cells (Fig. S6H) and specifically reduced the proportion of CD38^hi^ cells, without significantly affecting the CD38^dim^ subset (Fig. 7C-D). Given the role of CD38 as a major NAD⁺-consuming enzyme, we next quantified intracellular NAD⁺ levels in purified CD8⁺ T cells. CD8⁺ T cells from NR-treated mice displayed increased intracellular NAD⁺ levels relative to CD8⁺ T cells from PBS-treated infected animals (Fig. 7E).

**Figure 7.**
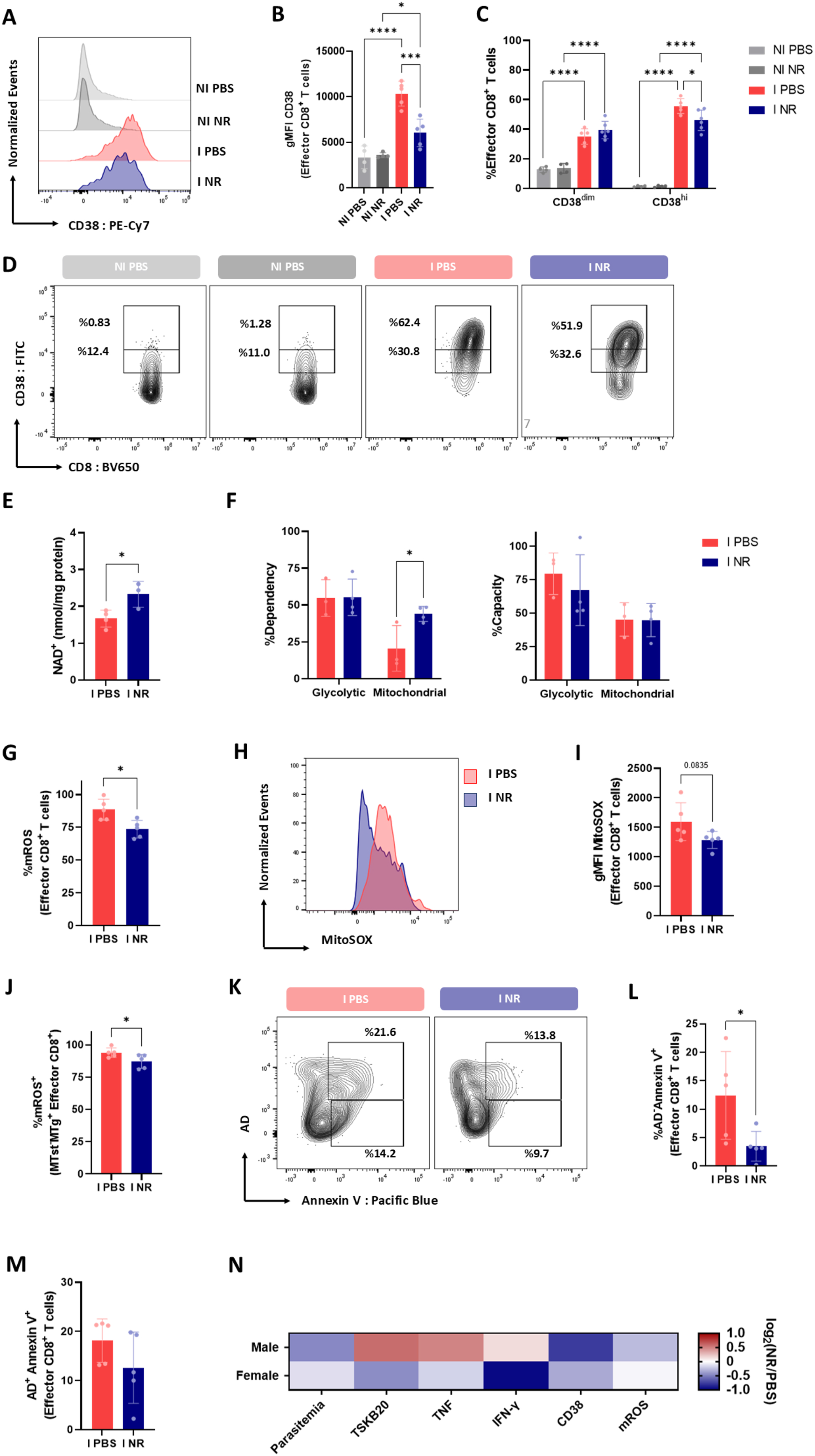
NR reduces CD38 expression and metabolic stress in CD8⁺ T cells during acute infection. Using the experimental design described in Figure 6, metabolic and phenotypic parameters were evaluated in effector CD8⁺ T cells. **(A-B)** CD38 expression was analyzed by flow cytometry. **(A)** Representative histograms of CD38 expression. **(B)** Mean fluorescence intensity (MFI) of CD38. **(C-D)** Effector CD8⁺ T cells were stratified into CD38^dim^ and CD38^hi^ populations. **(C)** Frequency of CD38^dim^ and CD38^hi^ cells. **(D)** Representative contour plots. **(E)** Intracellular NAD⁺ levels were quantified in purified CD8⁺ T cells by high-performance liquid chromatography (HPLC). **(F)** SCENITH analysis was performed on splenocytes stimulated with plate-bound anti-CD3 (5 μg/mL) and soluble anti-CD28 (1 μg/mL) for 18 h. Bar graphs show glycolytic dependence, mitochondrial dependence, glycolytic capacity, and mitochondrial capacity of effector CD8⁺ T cells. **(G-I)** Mitochondrial reactive oxygen species (mROS) were evaluated using MitoSOX. **(G)** Frequency of mROS⁺ effector CD8⁺ T cells. **(H)** Representative histograms of MitoSOX fluorescence. **(I)** Mean fluorescence intensity (MFI) of MitoSOX. **(J)** Frequency of mROS⁺ cells within the effector CD8⁺ T-cell population with depolarized mitochondria (MTst⁻MTg⁺). **(K-M)** Apoptosis was evaluated by Annexin V staining in combination with Aqua Dead (AD) viability dye. **(K)** Representative contour plots. **(L)** Frequency of AD⁻Annexin V⁺ effector CD8⁺ T cells. **(M)** Frequency of AD⁺Annexin V⁺ effector CD8⁺ T cells. **(N)** Heatmap showing the log₂ fold change (log₂[NR/PBS]) of the indicated parameters in male and female mice during the acute phase of *T. cruzi* infection. Values were calculated from the mean of each treatment group within each sex. Blue and red indicate decreases and increases, respectively, following NR supplementation, with white indicating no change relative to PBS-treated mice. TSKB20⁺ CD8⁺ T cells represent *T. cruzi*-specific CD8⁺ T cells. Data were obtained from I PBS (*n* = 5) and I NR (*n* = 5) mice for panels (B), (C), (G), (I), (J), (L), and (M); from I PBS (*n* = 4) and I NR (*n* = 3) mice for panel (E); and from I PBS (*n* = 3) and I NR (*n* = 4) mice for panel (F). For panel (N), data were obtained from male I PBS (*n* = 5), female I PBS (*n* = 4), male I NR (*n* = 5), and female I NR (*n* = 4) mice. Data are presented as mean ± SD. Data in (B) were analyzed using one-way ANOVA followed by Tukey’s multiple-comparisons test. Data in (C) and (F) were analyzed using two-way ANOVA followed by Šídák’s multiple-comparisons test. Data in (E), (G), (I), (J), (L), and (M) were analyzed using an unpaired two-tailed Student’s t-test. \**p* < 0.05, \*\*\**p* < 0.001, and \*\*\*\**p* < 0.0001.

To determine whether the increase in intracellular NAD⁺ levels was associated with changes in cellular metabolism, we next assessed the metabolic profile of effector CD8⁺ T cells using SCENITH. Effector CD8⁺ T cells from NR-treated mice exhibited increased mitochondrial dependency compared with cells from PBS-treated infected animals (Fig. 7F). By contrast, glycolytic dependency and glycolytic capacity remained unchanged between groups (Fig. 7F).

We next evaluated whether NR treatment modulated mitochondrial parameters in effector CD8⁺ T cells. NR-treated mice exhibited a reduced frequency of mROS^+^ cells, along with lower mROS levels compared with PBS-treated controls (Fig. 7G-I).

In contrast, NR treatment did not significantly affect mitochondrial mass (Fig. S6I) or the proportion of cells with depolarized mitochondria (Fig. S6J). Moreover, a reduced proportion of mROS⁺ cells was also observed within the population of cells with depolarized mitochondria in NR-treated mice (Fig. 7J).

Finally, NR-treated mice exhibited a reduced frequency of early apoptotic effector CD8⁺ T cells compared with PBS-treated infected animals (Fig. 7K-M).

We next assessed whether the effects of NR observed in male mice were similarly detected in females. In female mice, NR supplementation did not significantly alter the parameters evaluated, with the exception of the frequency of IFN-γ⁺ cells, which was reduced following NR treatment (Fig. S7A-G). Overall, the antiparasitic, immunological, and metabolic effects observed in male mice were largely absent in females (Fig. 7N).

Collectively, these findings indicate that NR supplementation in male mice elevated intracellular NAD⁺ levels and was associated with increased mitochondrial dependency, reduced oxidative stress, and decreased apoptosis in effector CD8⁺ T cells during acute infection.

### NR modulates memory CD8⁺ T cell phenotype and function during chronic *T. cruzi* infection

To determine whether the effects of NR treatment persisted beyond the acute phase, mice treated with NR during acute infection were analyzed at 112 days post-infection (dpi) (Fig. 8A). NR treatment did not significantly affect survival, body weight, or blood glucose levels during the chronic phase (Fig. 8B-D). At 112 dpi, NR-treated mice exhibited reduced parasite burden in the liver (Fig. 8E), whereas no differences were detected in skeletal muscle (Fig. S8A).

**Figure 8.**
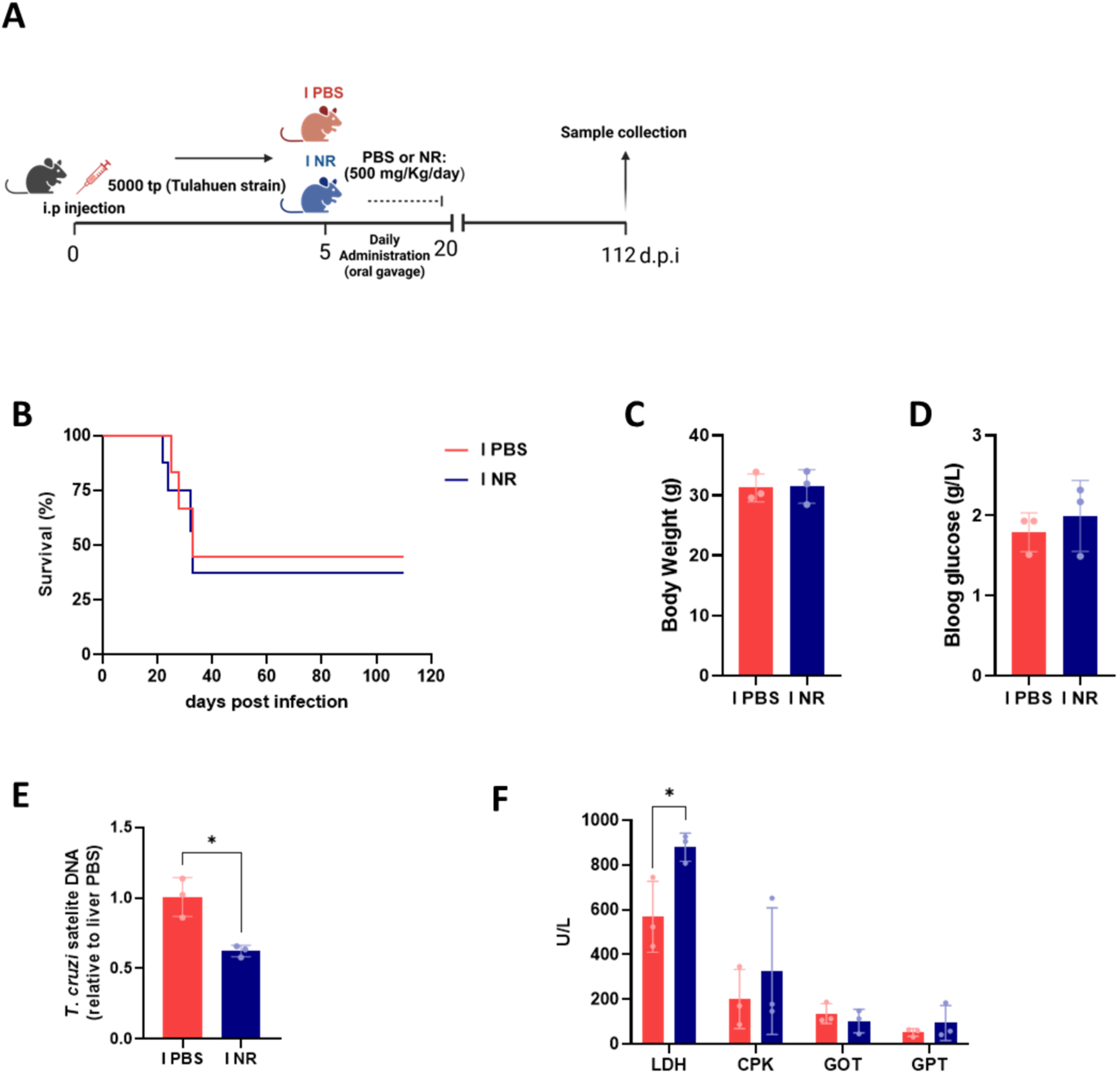
NR modulates memory CD8⁺ T cell phenotype and function during chronic *T. cruzi* infection. **(A)** Schematic representation of the experimental design. Male C57BL/6 mice were infected with 5,000 bloodstream trypomastigotes and treated daily with PBS or nicotinamide riboside (NR; 500 mg/kg) by oral gavage from 5 to 20 days post-infection (dpi). Mice were analyzed at 112 dpi. **(B)** Survival was monitored throughout the experimental period (0–112 dpi). **(C)** Body weight at 112 dpi. **(D)** Blood glucose levels at 112 dpi. **(E)** Liver parasite burden was quantified by real-time PCR in pooled liver samples from I PBS (*n* = 3) and I NR (*n* =3) mice and expressed relative to the I PBS group. **(F)** Plasma biochemical markers of tissue injury, including lactate dehydrogenase (LDH), creatine phosphokinase (CPK), glutamate oxaloacetate transaminase (GOT), and glutamate pyruvate transaminase (GPT). Data were obtained from I PBS (*n* = 3) and I NR (*n* = 3) mice. Data are presented as mean ± SD. Data in (C), (D) and (E) were analyzed using an unpaired two-tailed Student’s t-test. Data in (F) were analyzed using two-way ANOVA followed by Šídák’s multiple-comparisons test. \**p* < 0.05.

Moreover, serum markers of tissue damage, including GOT, GPT, and CPK, were comparable between groups, although an increase in LDH levels was observed in NR-treated mice (Fig. 8F).

We next characterized the memory CD8⁺ T cell compartment (CD44⁺CD62L⁺). NR-treated mice exhibited an increased frequency of total memory CD8⁺ T cells, whereas the frequency of parasite-specific memory CD8⁺ T cells remained unchanged (Fig. 9A-C). Phenotypic analysis revealed a reduction in the proportion of CD38^hi^ memory CD8⁺ T cells, with no significant changes in the CD38^dim^ subset (Fig. 9D-F). In addition, NR-treated mice displayed reduced expression levels of PD-1 and LAG-3 in memory CD8⁺ T cells (Fig. 9G).

**Figure 9.**
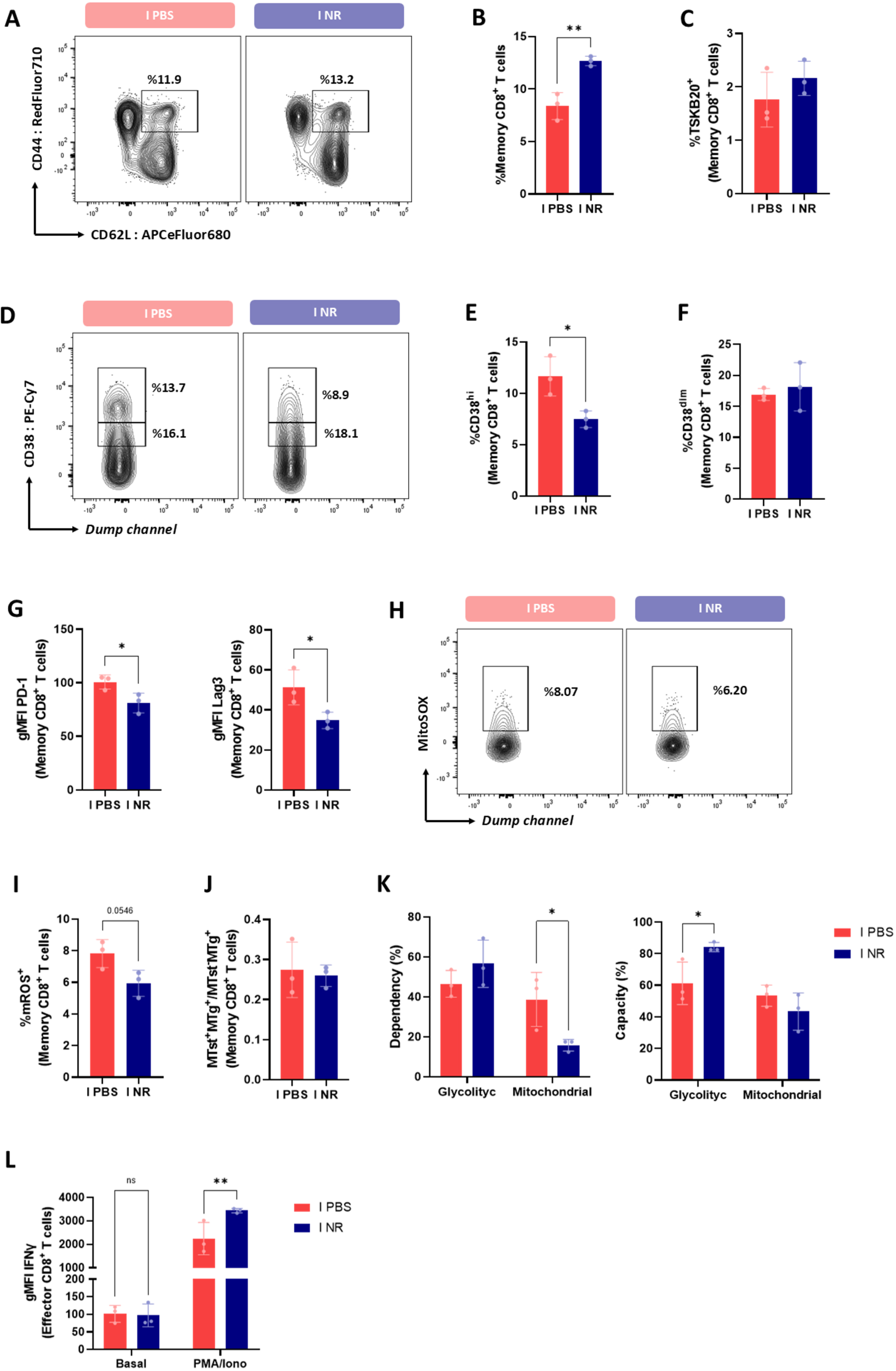
NR modulates memory CD8⁺ T cell phenotype and function during chronic *T. cruzi* infection. Memory CD8⁺ T cell populations were analyzed in the spleens of male C57BL/6 mice at 112 days post-infection (dpi) following treatment with PBS or nicotinamide riboside (NR). **(A)** Representative contour plots showing memory CD8⁺ T cells (CD44⁺CD62L⁺). **(B)** Frequency of memory CD8⁺ T cells. **(C)** Frequency of parasite-specific (TSKB20) cells within the memory CD8⁺ T-cell population. **(D-F)** CD38 expression was evaluated in memory CD8⁺ T cells by flow cytometry. **(D)** Representative contour plots showing CD38^dim^ and CD38^hi^ populations. **(E)** Frequency of CD38^hi^ memory CD8⁺ T cells. **(F)** Frequency of CD38^dim^ memory CD8⁺ T cells. **(G)** Mean fluorescence intensity (MFI) of PD-1 and LAG-3 in memory CD8⁺ T cells. **(H-I)** Mitochondrial reactive oxygen species (mROS) production was evaluated in memory CD8⁺ T cells using MitoSOX. **(H)** Representative contour plots. **(I)** Frequency of mROS⁺ memory CD8⁺ T cells. **(J)** MitoSpy™ Green (MTg), which labels mitochondria independently of mitochondrial membrane potential, together with the membrane potential-dependent probe MitoStatus Red (MTst), was used to identify memory CD8⁺ T cells with preserved mitochondrial membrane potential (MTst⁺MTg⁺) or depolarized mitochondria (MTst⁻MTg⁺). The ratio of MTst⁺MTg⁺ to MTst⁻MTg⁺ memory CD8⁺ T cells is shown. **(K)** SCENITH analysis was performed on splenocytes restimulated with plate-bound anti-CD3 (5 μg/mL) and soluble anti-CD28 (1 μg/mL) for 18 h. Bar graphs show glycolytic dependency, mitochondrial dependency, glycolytic capacity, and mitochondrial capacity of memory CD8⁺ T cells. **(L)** IFN-γ expression in memory CD8⁺ T cells following PMA/ionomycin stimulation, expressed as MFI. Data were obtained from I PBS (*n* = 3) and I NR (*n* = 3) mice and are presented as mean ± SD. Data in (B), (C), (E), (F), (G), (I), and (J) were analyzed using an unpaired two-tailed Student’s t-test. Data in (K) and (L) were analyzed using two-way ANOVA followed by Šídák’s multiple-comparisons test. \**p* < 0.05 and \*\**p* < 0.01.

In agreement with these findings, memory CD8⁺ T cells from NR-treated mice exhibited a reduced frequency of mROS^+^ cells (Fig. 9H-I), although no significant differences were observed in the proportion of cells with depolarized mitochondria (Fig. 9J).

Metabolic profiling by SCENITH revealed reduced mitochondrial dependency together with increased glycolytic capacity in memory CD8⁺ T cells from NR-treated mice after polyclonal stimulation (Fig. 9K). Importantly, memory CD8⁺ T cells from NR-treated mice showed increased IFN-γ production upon stimulation (Fig. 9L).

Together, these findings indicate that transient NR supplementation during acute infection was associated with long-lasting phenotypic, metabolic, and functional remodeling of the memory CD8⁺ T cell compartment, which was detectable into the chronic phase.

## Discussion

T cell metabolic reprogramming is a key determinant of T cell differentiation and function (39). While aerobic glycolysis supports rapid proliferation and effector differentiation (13, 40), mitochondrial metabolism is now recognized as a critical regulator of CD8⁺ T cell proliferation, differentiation, and long-term functional fitness (15, 17, 41). Therefore, disruption of mitochondrial metabolic homeostasis may compromise the quality of CD8⁺ T cell responses during infection.

In this context, our findings show that the metabolic changes associated with the CD8⁺ T cell response during acute *T. cruzi* infection are accompanied by mitochondrial stress. Accumulating evidence identifies the CD38-NAD⁺ axis as an important regulator of T cell immunometabolism. Experimental studies have demonstrated that sustained CD38 expression promotes intracellular NAD⁺ depletion, leading to impaired SIRT1 signaling, mitochondrial dysfunction, increased oxidative stress, and defective effector responses in CD8⁺ T cells (22, 42). Consistent with these mechanisms, elevated CD38 expression in cytomegalovirus-specific CD8⁺ T cells has been associated with mitochondrial stress and metabolic dysfunction, which can be partially reversed by inhibition of CD38 (43).

Our findings are consistent with this model and suggest that the CD38-NAD⁺ axis may be involved in the metabolic remodeling of CD8⁺ T cells during acute *T. cruzi* infection. Acute infection was associated with the accumulation of CD38^hi^ cells within both parasite-specific and effector CD8⁺ T cell populations, and this subset displayed a phenotype characterized by mitochondrial alterations and increased apoptosis. Reduced SIRT1 expression together with the trend toward lower intracellular NAD⁺ levels observed in CD8⁺ T cells from infected mice are consistent with this interpretation. Although intracellular NAD⁺ was quantified in total CD8⁺ T cells rather than in CD38-defined subsets, recent studies have demonstrated that CD38^hi^ T cells exhibit lower intracellular NAD⁺ levels together with impaired mitochondrial fitness and diminished cytotoxic function than their CD38^low^ counterparts (42, 44). Both genetic ablation and pharmacological inhibition of CD38 restore intracellular NAD⁺ homeostasis, mitochondrial function, and T cell effector function, further supporting a potential role for CD38-dependent NAD⁺ metabolism in shaping T cell metabolic fitness (42, 44).

Maintenance of NAD⁺ homeostasis is increasingly recognized as a key determinant of immune cell metabolism and function (21, 23). Accordingly, strategies that increase intracellular NAD⁺ availability have been explored to improve mitochondrial fitness and T cell function. Restoration of intracellular NAD⁺ levels improved mitochondrial fitness in aged CAR-T cells (25), maintenance of de novo NAD⁺ synthesis preserved the metabolic programs required for antitumor CD8⁺ T cell responses (24), and NR supplementation enhanced mitochondrial fitness and T cell function in tumor models (27, 45).

Interestingly, beyond serving as an NAD⁺ precursor, NR has recently been identified as a covalent inhibitor of CD38 NADase activity, suggesting an additional potential mechanism through which NR may influence intracellular NAD⁺ homeostasis (46). However, CD38 NADase activity was not directly assessed in the present study, and the mechanism underlying the reduced CD38 expression observed following NR treatment remains unknown. Thus, the relative contribution of enhanced NAD⁺ biosynthesis, CD38 NADase inhibition, and reduced CD38 expression to the immunometabolic effects observed in the present study remains to be determined. Consistent with these observations, NR supplementation increased intracellular NAD⁺ levels and was associated with improved CD8⁺ T cell metabolic fitness and effector responses *in vivo*. These immunometabolic changes were accompanied by improved parasite control during acute infection, without overt changes in the measured systemic markers of tissue injury or disease severity.

Moreover, although NR may influence multiple immune and non-immune cell populations *in vivo* (23), our *in vitro* differentiation experiments support direct effects on CD8⁺ T cells. Differentiation of CD8⁺ T cells in the presence of NR enhanced mitochondrial metabolism, reduced oxidative stress, and promoted proliferation and cytokine production without altering effector differentiation. These observations are consistent with a recent *in vitro* model of CD8⁺ T cell exhaustion, in which NR alleviated mitochondrial dysfunction, reduced apoptosis, and preserved T cell function through a SIRT1-dependent mechanism (47). Nevertheless, we cannot exclude that modulation of additional immune or non-immune cell populations also contributed to the improved parasite control observed *in vivo* and the direct contribution of CD8⁺ T cells to this effect remains to be determined. Overall, improved parasite control is unlikely to reflect a direct trypanocidal effect of NR. Although NR exhibited some direct antiparasitic activity *in vitro*, particularly at the highest concentration tested, its activity remained substantially lower than that of benznidazole (Fig. S8B). These findings suggest that the improved parasite control observed *in vivo* may involve modulation of host immune responses, including the CD8⁺ T cell response, rather than being explained solely by direct antiparasitic activity.

Beyond its effects during acute infection, transient NR administration during the acute phase increased the frequency of memory CD8⁺ T cells. These cells exhibited a phenotype characterized by reduced CD38, PD-1, and LAG-3 expression, decreased mitochondrial oxidative stress, and enhanced IFN-γ production upon restimulation. Given that memory CD8⁺ T cell differentiation and maintenance are tightly linked to metabolic programming (48), these findings suggest that transient NR supplementation during acute infection is associated with persistent phenotypic, functional, and metabolic remodeling of memory CD8⁺ T cells. This interpretation is consistent with recent evidence showing that NR promotes memory differentiation together with improved metabolic fitness and recall responses of CD8⁺ T cells during adoptive cell therapy (45). Nevertheless, the beneficial effects of NR during chronic infection were only partial. Although NR-treated mice exhibited reduced parasite burden in the liver, parasite load remained unchanged in skeletal muscle. These findings underscore the multifactorial nature of chronic *T. cruzi* persistence and highlight the limited impact of transient NR supplementation on parasite control during chronic infection. Future studies should determine whether prolonged NR administration or combination with trypanocidal drugs could enhance long-term parasite control, particularly given the reported safety and tolerability of NR in humans and its oral bioavailability (49, 50).

Notably, the metabolic and immunological effects of NR observed in male mice were largely not detected in females. These sex-dependent differences may reflect, at least in part, previously described sexual dimorphism in the host response to *T. cruzi* infection. These studies have reported higher parasitemia, mortality, and severity of myocarditis in infected male mice compared with females, although the magnitude of these differences can vary depending on both host genetic background and parasite strain (35). Sex hormones have also been implicated in this differential susceptibility, with testosterone promoting, and estradiol limiting, disease severity during acute infection (34). Thus, the distinct response to NR observed between male and female mice may reflect, at least in part, underlying sex-dependent differences in the immune, metabolic, and hormonal environment established during *T. cruzi* infection. More broadly, sex-dependent differences in energy metabolism and immune function are increasingly recognized (51, 52), and sexually dimorphic responses to interventions targeting the CD38-NAD⁺ axis have also been reported (53). These observations underscore the importance of considering sex as a biological variable when investigating immunometabolic interventions and suggest that response to NAD⁺-boosting strategies may depend on the metabolic and hormonal context of the host.

Taken together, our findings provide evidence that acute *T. cruzi* infection is associated with metabolic and mitochondrial alterations in parasite-specific and effector CD8⁺ T cells, accompanied by increased CD38 expression. NR supplementation increased intracellular NAD⁺ levels, reduced metabolic stress, improved mitochondrial fitness, enhanced CD8⁺ T cell effector responses, and was associated with improved parasite control. Collectively, these findings provide a rationale for further investigating NAD⁺-boosting strategies as host-directed immunometabolic interventions against *T. cruzi* infection.

## Materials and Methods

Key reagents and resources used throughout the study, including their sources and identifiers, are listed in Table S1.

### Mice

Male and female BALB/c and C57BL/6 mice were obtained from the School of Veterinary Sciences, La Plata National University (La Plata, Argentina). Animals were bred in the animal facility of the Faculty of Chemical Sciences, National University of Córdoba and housed under a 12:12 h light-dark cycle with food and water ad libitum. All animal procedures were approved by the Animal Care and Use Committee of the Faculty of Chemical Sciences, National University of Córdoba (Approval Numbers HCD 1402/22 and 137/26) and conducted in accordance with institutional guidelines for the care and use of laboratory animals.

### Mouse model of *T. cruzi* infection

*T. cruzi* Tulahuen strain bloodstream trypomastigotes were maintained in male BALB/c mice by serial passages every 10–11 days (54).

For experimental infections, C57BL/6 mice, approximately 8 weeks old, were intraperitoneally infected with 5,000 blood-derived *T. cruzi* trypomastigotes. Male mice were used for characterization of infection-induced CD8⁺ T cell responses and metabolic remodeling. For *in vivo* nicotinamide riboside (NR) treatment experiments, both male and female mice were included and analyzed separately. Mice were analyzed during the acute phase (20–21 days post-infection, dpi) and the chronic phase (112 dpi). Mice were anesthetized with isoflurane. Animals were euthanized by cervical dislocation, and tissues were collected for subsequent analyses. Age-matched non-infected (NI) mice were used as controls.

### Nicotinamide Riboside treatment

Male and female mice were treated daily with NR chloride (500 mg/kg/day, Renue by Science) by oral gavage in 100 μL of PBS from 5 to 20 dpi. Control mice received PBS alone.

### Parasite quantification in blood and tissues

Parasitemia was assessed by counting viable trypomastigotes in a Neubauer chamber after red blood cell lysis. For tissue parasite quantification, genomic DNA was purified from 50 μg of tissue using TRIzol following the manufactureŕs instructions. Satellite DNA from *T. cruzi* was quantified by real time PCR using previously described primer and probe sequences (55). *T. cruzi* satellite DNA levels were normalized to GAPDH using the comparative Ct method and expressed as arbitrary units, as previously reported (56).

### Biochemical determinations in plasma

Plasma levels of lactate dehydrogenase (LDH), creatine phosphokinase (CPK), glutamate pyruvate transaminase (GPT), and glutamate oxaloacetate transaminase (GOT) were quantified at Biocon Laboratory (Córdoba, Argentina).

### Isolation of splenocytes and CD8^+^ T cell purification

Splenocytes were obtained by mechanical disaggregation and red blood cell lysis and resuspended in RPMI containing 10% FBS, 2 mM L-glutamine, and 40 µg/mL gentamicin. CD8⁺ T cells were purified by negative selection using the MojoSort Mouse CD8 T Cell Isolation Kit yielding 95–98% purity.

### Flow cytometry

Unless otherwise indicated, 5 × 10⁶ cells were used for each staining condition. Cell suspensions were washed in PBS supplemented with 2% FBS and incubated with LIVE/DEAD Fixable Aqua Dead Cell Stain Kit together with fluorochrome-conjugated antibodies for surface staining for 20 min at 4 °C. For identification of *T. cruzi*-specific CD8⁺ T cells, cells were incubated with the H-2K^b^ TSKB20 (ANYKFTLV) Brilliant Violet 421-labeled tetramer for 20 min at 4 °C together with surface antibodies.

To assess mitochondrial reactive oxygen species (mROS) levels, cells were incubated for 15 min at room temperature with 5 μM MitoSOX™ Red.

For analysis of mitochondrial parameters, cells were incubated for 20 min at 37 °C with MitoSpy™ Green FM (100 nM) to assess mitochondrial mass, together with either MitoStatus Red (100 nM) or TMRE (50 nM) to assess mitochondrial membrane potential.

For glucose uptake assessment, 2×10^6^ cells per condition were cultured in low-glucose Dulbecco’s Modified Eagle Medium (DMEM) supplemented with 1% FBS for 30 min, followed by incubation with 10 µM 2-NBDG for 40 min at 37 °C.

For transcription factor detection, cells were stained for surface markers, washed, fixed, permeabilized and stained with Foxp3/Transcription Factor Staining Buffers. For intracellular cytokine detection, 2 ×10^6^ cells per well were cultured in 200 μL supplemented RPMI 1640 medium and stimulated for 5 h at 37 °C with 50 ng/mL PMA and 1 μg/mL ionomycin in the presence of Brefeldin A and Monensin. Stimulated cells were then surface-stained, fixed, and permeabilized using the BD Cytofix/Cytoperm™ Fixation/Permeabilization Kit.

Metabolic profiling of CD8⁺ T cells was performed using the SCENITH (Single-Cell ENergetIc metabolism by Translation inHibition) assay as previously described (37). Splenocytes isolated from non-infected (NI) and infected mice were seeded in 48-well plates at a density of 2 × 10⁶ and 5 × 10⁵ cells per well, respectively, and stimulated for 16–18 h with plate-bound anti-CD3 and soluble anti-CD28, or *T. cruzi* lysates (8.34 μg/mL). Following stimulation, cells were harvested, washed, and transferred to 96-well plates, where they were incubated with the metabolic inhibitors 2-deoxy-D-glucose (2-DG), oligomycin (O), or both (DGO). Cells incubated without metabolic inhibitors served as controls (Co). Puromycin was then added to label newly synthesized proteins. Cells were subsequently stained for viability and surface markers, fixed, permeabilized, and stained intracellularly with Alexa Fluor 647-conjugated anti-puromycin antibody. Metabolic parameters, including glycolytic dependency, mitochondrial dependency, glycolytic capacity, and mitochondrial capacity, were calculated as previously described (37). Intracellular staining was performed for 1 h at 4° C.

Samples were acquired on a CytoFLEX S (Beckman Coulter), BD LSRFortessa (BD Biosciences) or Northern Lights™ spectral flow cytometer (Cytek), and data were analyzed using FlowJo software (v10; Becton, Dickinson and Company, Ashland, OR, USA).

### Measurement of NAD^+^ levels by high-performance liquid chromatography (HPLC)

Intracellular NAD⁺ levels in purified CD8⁺ T cells were determined by high-performance liquid chromatography (HPLC) as previously described (57). Briefly, cell pellets were resuspended in 0.05 M phosphate buffer (pH 7.0; buffer A) and deproteinized with 30% (w/v) trichloroacetic acid (TCA) at a 4:1 (v/v) ratio of buffer A to TCA. Then, 40 μL of each sample was injected into an Agilent 1220 HPLC system equipped with a binary pump and a variable-wavelength UV detector set at 261 nm. The total run time was 30 min. Data acquisition and analysis were performed using Agilent OpenLAB CDS software. Chromatographic separation was achieved on a Phenomenex® C18 reverse-phase column (150 × 4.6 mm, 5 μm particle size). The mobile phase consisted of 0.05 M phosphate buffer, pH 7.0 (A) and methanol (B) at a flow rate of 1 mL/min. The elution program was as follows: 100% A from 0–5 min; a linear gradient to 95% A/5% B from 5–6 min; 95% A/5% B from 6–11 min; a linear gradient to 85% A/15% B from 11–13 min; 85% A/15% B from 13–23 min; a linear gradient back to 100% A from 23–24 min; and 100% A from 24–30 min. Peak identity was confirmed by co-injection with an NAD⁺ standard. Quantification was performed by comparing sample peak areas with a calibration curve generated using an NAD⁺ standard, and NAD⁺ levels were normalized to total protein content determined by Bradford assay.

### Generation of effector CD8⁺ T cells *in vitro*

Naïve CD8⁺ T cells were isolated using the MojoSort Mouse Naïve CD8 T Cell Isolation Kit. Cells were differentiated toward an effector phenotype in the presence of anti-CD3 (10 µg/mL), anti-CD28 (1 µg/mL), anti-IL-4 (10 µg/mL), and recombinant mouse IL-12 (20 ng/mL). Cells were cultured in RPMI 1640 containing 2 mM GlutaMAX, supplemented with 10% FBS and 1% penicillin/streptomycin. NR was added to the culture medium at a final concentration of 40 μM throughout the differentiation period. Control cultures received an equivalent volume of DMSO to match the final vehicle concentration.

### Cell proliferation

For proliferation assays, naïve CD8⁺ T cells were labeled with CellTrace™ Violet for 7 min at 37 °C in the dark, according to the manufacturer’s instructions. Cells were then washed and plated at 1 × 10⁵ cells per well in 96-well plates under the differentiation conditions described above. At the end of the culture period, cells were stained with LIVE/DEAD™ Fixable Aqua viability dye and surface antibodies. Proliferation was evaluated by CellTrace Violet dilution within the viable CD8⁺ T cell population.

### Seahorse metabolic analysis

*In vitro*-differentiated effector CD8⁺ T cells (1.5 × 10⁵ cells per well) were attached to Seahorse XFe96 cell culture microplates coated with 0.01% poly-D-lysine. Metabolic analyses were performed using a Seahorse XFe96 Extracellular Flux Analyzer (Agilent Technologies, USA).

For the mitochondrial stress test, culture medium was replaced with Seahorse XF Base Medium (pH 7.4) supplemented with 2 mM L-glutamine, 5 mM glucose, and 1 mM sodium pyruvate. Oligomycin (5 μM), carbonyl cyanide-p-trifluoromethoxyphenylhydrazone (FCCP, 1 μM), and a mixture of antimycin A (AA, 5 μM) and rotenone (Rot, 1 μM) were sequentially injected. Oxygen consumption rate (OCR) was measured, and basal respiration, maximal respiration, ATP-linked respiration, and non-mitochondrial respiration were calculated.

For the glycolytic rate assay, cells were plated in Seahorse XF Glycolytic Rate Assay Medium supplemented with 5 mM glucose and 10 mM HEPES (pH 7.4). Basal proton efflux rate (PER) was recorded. Subsequently, AA (5 μM) and Rot (1 μM) were injected. Glycolytic proton efflux rate (glycoPER), basal glycolysis, and compensatory glycolysis were calculated.

### Gene expression analysis by RT-qPCR

Total RNA was isolated from 1 - 2 × 10⁶ CD8⁺ T cells differentiated *in vitro* using TRIzol reagent, followed by purification with the Direct-zol RNA Microprep Kit. RNA concentration and purity were determined using a NanoDrop One spectrophotometer, and only samples with an A260/280 ratio between 1.8 and 2.3 were used for downstream analyses. cDNA was synthesized from 200 ng of total RNA using oligo(dT) primers and SuperScript III Reverse Transcriptase according to the manufacturer’s instructions. Quantitative real-time PCR was performed using gene-specific primers, and relative gene expression was calculated using the 2^−^ΔΔCt method with β-actin (Actb) as the reference gene.

### *In vitro* treatment of *T. cruzi* trypomastigotes

Blood-derived trypomastigotes were used to infect confluent monolayers of Vero cells (54). After 7 days, culture supernatants containing trypomastigotes were collected, and parasites were counted using a Neubauer chamber. A total of 3 × 10⁵ parasites were added to each well of a 96-well plate and incubated with 20, 40, or 80 µM NR diluted in RPMI 1640 medium for 24 h at 37 °C. Benznidazole (BZ) was used as a positive control, whereas parasites incubated with an equivalent volume of DMSO served as vehicle-treated controls.

### Body weight and survival evaluation

Body weight was determined using a precision laboratory balance. Survival of treated mice was monitored daily until the end of the experiment (21 or 112 dpi).

### Histological analysis

Liver and skeletal muscle were obtained from perfused animals, fixed in formaldehyde solution and embedded in paraffin. Five-μm-thick sections were stained with hematoxylin followed by eosin. Histopathological evaluation was performed by a pathologist under light microscopy. Images were acquired using a Phenoimager Fusion instrument (Akoya Biosciences, RRID:SCR_023274).

### Data Analysis

Statistical analyses were performed using GraphPad Prism version 9 (GraphPad Software, San Diego, CA, USA). Data distribution was assessed using the Shapiro–Wilk test. Differences between two groups were analyzed using a two-tailed Student’s t-test. Multiple groups or experimental factors were analyzed by one-way or two-way ANOVA, including repeated-measures designs when appropriate, Tukey’s or Šídák’s multiple-comparisons test, as appropriate. A *P* value < 0.05 was considered statistically significant. Data are presented as mean ± SD. Statistical tests and replicate numbers are specified in the figure legends.

Parasitemia, biochemical parameters, and histopathology outcomes were assessed by investigators blinded to treatment allocation.

## Acknowledgments

We thank the staff of our core facilities for their invaluable support: P. Abadie, S. Boccardo and P. Crespo for flow cytometry and microscopy, G. Furlan, N. Maldonado, and A. Romero for cell culture assistance, S. Oms, D. Lutti, V. Blanco, R. Villareal and C. Noriega for animal care, MS. Miró and D. Paira for histological procedures. CCS, MGT, CCM, FMC and EVAR are members of the Scientific Career of CONICET. FH, CF, REB, CMSG, MH thank CONICET for the fellowships granted. We acknowledge the NIH Tetramer Core Facility for provision of TSKB20 tetramers. We thank Dr. Arguello for kindly providing the SCENITH kit used in this study. We also thank Dr. Susana Genti for kindly providing the NAD⁺ and NADH standards used for HPLC analysis. The authors are grateful for the artwork (featured image) by David Rojas Márquez, scientific illustrator and PhD (GraficAr SCIENTIA).

## CRediT authorship contribution statement

Florencia Hellriegel: Conceptualization; Investigation; Methodology; Formal analysis, Validation; Visualization; Writing – original draft; Writing – review & editing. Camila Fontanari: Investigation; Methodology. Ruth E. Baigorri: Investigation; Methodology. Martin G. Theumer: Investigation, Methodology; Validation. Camila M. S. Gimenez: Investigation; Methodology. Melisa Herrera: Investigation; Methodology. Luciana Mezzano: Methodology; Formal analysis. Claudia C. Motrán: resources; Writing – review & editing. Fabio M. Cerban: Resources; Writing – review & editing. Eva V. Acosta Rodriguez: Methodology; Resources, Writing – review & editing. Luciana Berod: Resources; Supervision; Writing – review & editing. Cinthia C. Stempin: Conceptualization; Investigation; Formal Analysis; Supervision; Funding acquisition; Project administration; Writing – original draft, Writing – review & editing.

## Declaration of generative AI and AI-assisted technologies in the writing process

During the preparation of this work the authors used *ChatGPT (OpenAI)* in order to improve the clarity and grammar of the manuscript. After using this tool, the author(s) reviewed and edited the content as needed and take full responsibility for the content of the published article.

---

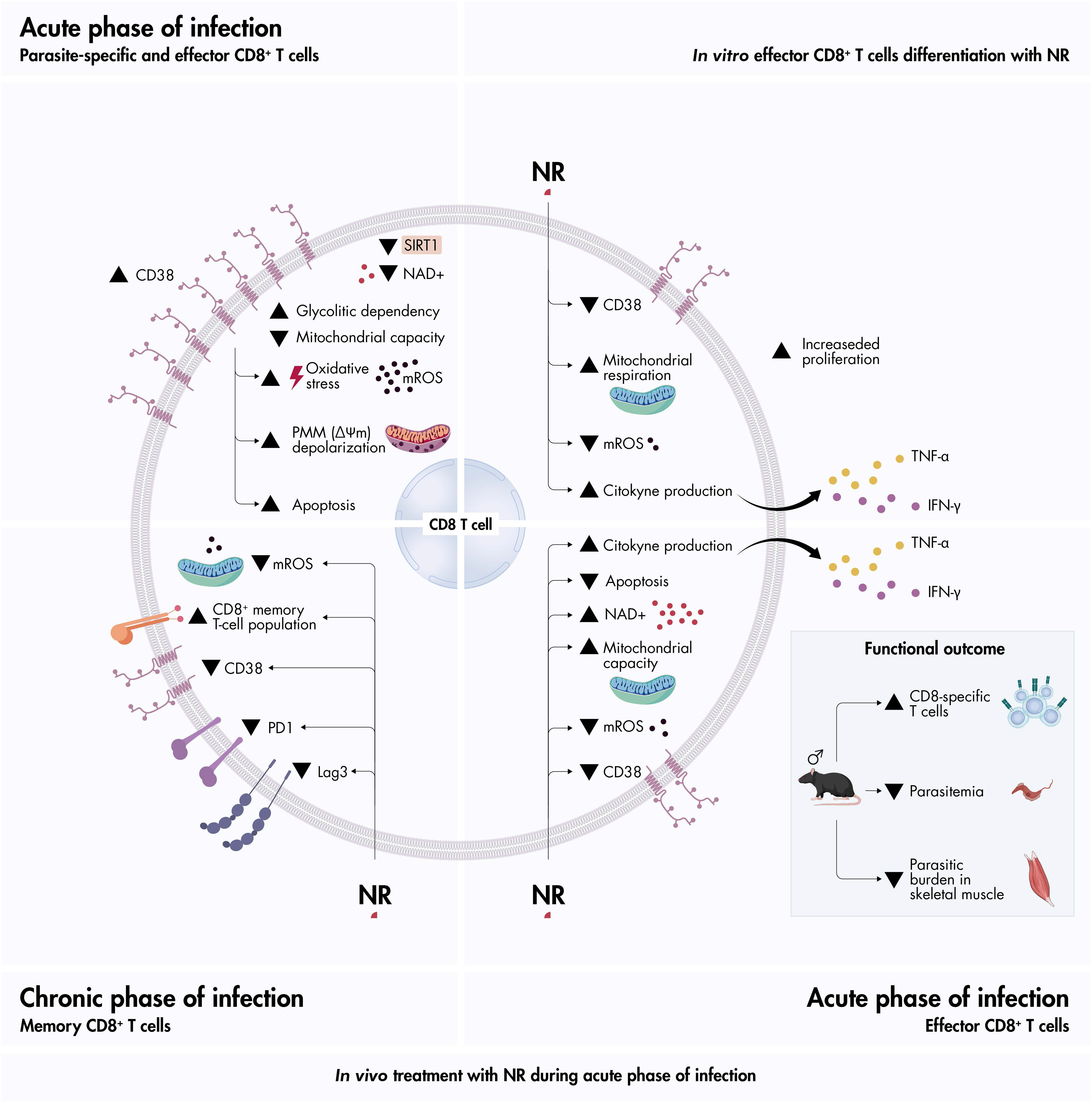

